# Recurrent inhibition crosses the spinal cord midline in humans

**DOI:** 10.64898/2026.08.18.745608

**Authors:** Julian Colard, Dorian Glories, Stéphane Baudry

## Abstract

Recurrent inhibition is known to modulate motoneuron output within an active motor pool, but it is unclear whether Renshaw cells receive projections from a contralateral pathway.

Using intramuscular single motor unit recordings in humans, we demonstrated that electrical activation of the contralateral quadriceps motor axons elicits a robust decrease in soleus motor unit discharge rate, consistent with the characteristic features of recurrent inhibition. The duration of the inhibition scaled with motor unit firing rates and exhibited substantial interindividual variability.

To uncover the underlying circuitry, we developed a biophysically grounded spiking network model constrained by individual experimental data. The model reproduced the observed contralateral inhibitory dynamics only when incorporating a polysynaptic commissural pathway mediated by V3-like interneurons. Model-based inference further revealed that intrinsic motoneuron properties critically shape the duration of inhibition. Together, these findings provide the first evidence for a commissural pathway influencing human spinal recurrent inhibitory networks, revealing a previously unrecognized mechanism that may contribute to bilateral motor coordination.

**Teaser:** A commissural spinal circuit contributes to recurrent inhibition of the soleus motor unit pool in humans.

## Introduction

During human movement, lower motoneuron (MN) activity is controlled through descending and peripheral inputs that selectively adjust motor output. Among these pathways, recurrent inhibition (RI) mediated by Renshaw cells (RC) is known to shape MN temporal dynamics (*1*). Recurrent collaterals from motor axons activate RC, which in turn form inhibitory synapses with MNs (*2*, *3*). Most MNs nuclei receive RI from their own collaterals (homonymous RI) and from MNs supplying other muscles acting synergistically at the same or at a different joint (heteronymous RI) (*3–5*). Thus, this inhibitory control extends beyond one motor pool, supporting a sophisticated role for RC in coordinating muscle activity. RI pathways are more extensively distributed in the human lower limb (*5*) than in hindlimbs of quadrupedal animals (*3*, *6*, *7*), likely reflecting phylogenetic adaptations to support bipedal stance and gait (*8*, *9*). RI limits temporal and spatial synchronization, and adjusts motor gain (*8*, *10*). Importantly, the strength of RI is not fixed and can be modulated by the level of motor activity (*8*, *11*).

RI can be assessed in humans with a method (M-only) originally developed by Kudina and Pantseva (*12*), and recently refined (*19*). The M-only method assesses RI by selectively stimulating motor axons with intensities that evoke a small M response, but no or very small H-reflex, which inhibits voluntarily firing a single motor unit (MU). Because this stimulation generates an antidromic action potential propagating centrally along the motoneuron axon toward the spinal cord, the action potential can invade recurrent axon collaterals and orthodromically activate RC within the spinal cord. This allows the resulting inhibition to be quantified through precise changes in single motor unit discharge rates (i.e., PSTH/PSF analysis). This approach provides greater validity than the classical paired H-reflex method, which activates mixed nerves and introduces major confounds such as Ia and Ib afferent activation, presynaptic inhibition and post-activation depression (*14*). To further reduce bias associated with homonymous stimulation paradigms (e.g., tibial nerve stimulation to assess RI in the soleus), which may be contaminated by afferent inputs and local circuit interactions, it is possible to use a heteronymous approach that avoids direct activation of the target motor pool, and thereby reduces these confounds and facilitates the dissociation of input and output pathways. Therefore, M-only heteronymous stimulation may isolate RI more specifically, enabling reliable estimation of its latency and duration, which are relevant to characterise the functioning of RI.

Growing evidence indicates that neural networks can have robust contralateral projections likely mediated by commissural interneurons (*15*). Human studies have shown that stimulation of afferents from the femoral nerve of one limb induces rapid inhibitory effects on contralateral soleus Hoffmann (H) reflex through an oligosynaptic commissural inhibitory pathway (*16*). Human studies also show that stimulation of contralateral posterior tibial and ipsilateral fibular nerve afferents converge onto shared Ia inhibitory interneurons within the reciprocal inhibition pathway, likely mediated by commissural interneurons that cross the midline (*17*). Additional work in humans and other animal models have shown that commissural interneurons, including V0 and V3 populations, contribute to crossed inhibitory reflexes and bilateral coordination during rhythmic locomotor tasks (*15*, *18*, *19*). Although ipsilateral RI is well characterized, it remains unknown whether this mechanism extends across the spinal midline to influence contralateral MN pools. Given RI’s prominent role in shaping motor output, such an organization would provide a particularly effective substrate for bilateral coordination, as reported for other pathways. V3 commissural interneurons, which project both ipsilaterally and contralaterally (*19*), are plausible candidates for relaying contralateral recurrent signals. However, while V3 interneurons have been shown to form substantial excitatory input onto spinal interneurons in mice, including RC (*19*), no direct evidence currently supports the functional existence of such a pathway, and their identification in the human spinal cord remains entirely unexplored. Therefore, we aimed to determine whether a commissural pathway, potentially involving V3-like interneurons, can relay recurrent inhibitory signals from one limb to the contralateral soleus motor unit pool.

To this end, we recorded single motor units from the soleus muscle conditioned by both ipsilateral and contralateral femoral nerve stimulation. We measured changes in MU discharge rate in response to ipsi– and contralateral femoral nerve stimulation with the rationale that ipsilateral RI provides reference values to be compared with those obtained from contralateral femoral nerve stimulation. Even though the M-only technique was designed to assess ipsilateral RI, we assumed this method can remain valid for assessing contralateral RI. Although intramuscular electromyography (iEMG) enables high-resolution recordings at the level of single motor units, its capacity to support mechanistic inference is inherently limited when used in isolation due to the restricted sampling of motor units. Therefore, to support experimental data, we used a spiking neural network model to determine whether the observed inhibitory signatures could emerge from a polysynaptic commissural pathway involving V3-like interneurons. In parallel, we examined how intrinsic MN properties, including soma capacitance, shape the strength of this contralateral pathway. Our results provide a framework for identifying a commissural RI circuit in the human spinal cord and reveal a potential mechanism supporting bilateral motor coordination.

## Results

### Characterization of ipsilateral heteronymous recurrent inhibition

In the ipsilateral condition, 62 motor units were recorded, of which 54 (87%) exhibited robust inhibition and were therefore retained for subsequent analyses. Analysis revealed a clear heteronymous ipsilateral RI in the soleus muscle (Figure 1A) with a mean inhibition latency of 21.5 ± 2.3 ms and an average duration of 35.6 ± 7.4 ms. Because RI properties might depend on motor unit firing rate, we examined the linear relationship between firing rate and inhibition duration, as well as firing rate and inhibition latency (*20*, *21*). We used the coefficient of determination (r²) to quantify these relationships. As shown in Figure 1B, when we pooled all motor units across participants, firing rate only moderately contributed to inhibition duration in the ipsilateral condition (r² = 0.35). These results are consistent with previous studies employing similar pooled analytical approaches (*22*). Notably, linear extrapolation of inhibition duration to rest predicted a value of 46.9 ms. Because interindividual variability substantially contributes to these relationships, we fitted linear models separately for each participant (LM_exp_). In the ipsilateral condition (Figure 1C), three participants (S4, S6, and S8) exhibited a strong negative relationship (r² > 0.67). Four participants showed moderate relationships (0.33 < r² < 0.67; S2, S5, S12, and S13), while the remaining six participants showed weak or absent relationships (r² < 0.33).

**Figure 1.**
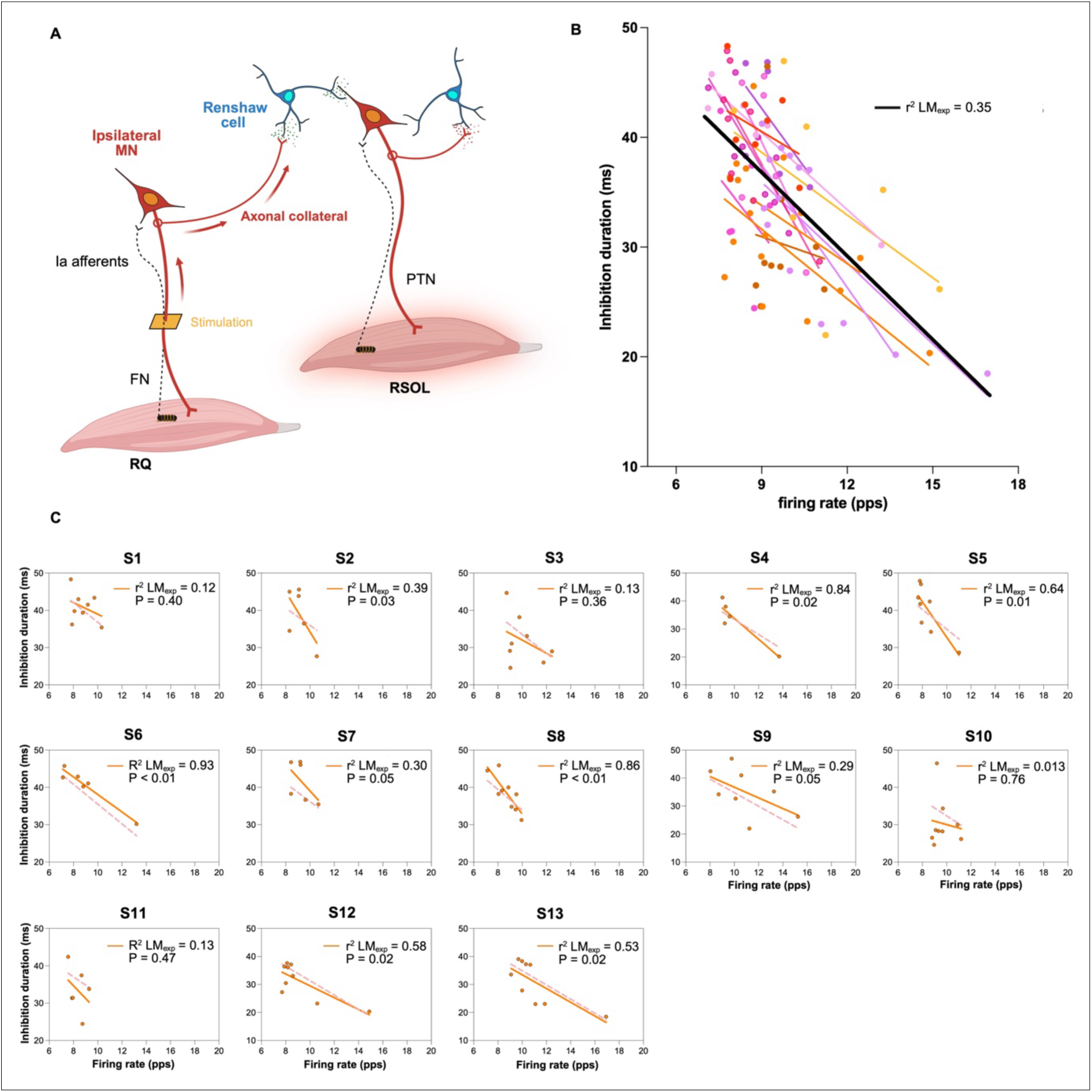
Characterisation of ipsilateral heteronymous recurrent inhibition duration. A) Schematic representation of the ipsilateral recurrent inhibition pathway (red). Electrical stimulation of right quadriceps motor axons was delivered at a constant intensity (approximately 10% of the maximal M-wave) and at intervals between 1 and 2 s, activating MN recurrent collaterals. These collaterals synapse onto Renshaw cells within the spinal cord, which in turn project back onto MNs innervating the right soleus muscle. B) Estimation of Ipsilateral recurrent inhibition duration of single MUs and their firing rate across participants. Colored dots represent MUs and the linear regression fit per individual; black line represents the linear regression fit for the group data with respective r² shown. (C) Inhibition duration of individual motor units (MUs) as a function of firing rate for each participant (orange dots). Orange lines represent individualized linear fits (LM_exp_) with the corresponding r² and P values, whereas pink dotted lines represent the linear mixed-effects model (LMM) regression for each participant. MN, motoneuron; FN, femoral nerve; PTN, posterior tibial nerve; RQ, right quadriceps, RSOL, right soleus.

Regarding inhibition latency, our pooled analysis revealed no relationship with motor unit firing rate in the ipsilateral condition (r² = 0.00). Consistently, no participant exhibited a significant relationship at the individual level (all r² < 0.19). These results demonstrate that, unlike inhibition duration, the latency of ipsilateral heteronymous RI remains independent of motor unit firing rate, suggesting a constant conduction delay within the recurrent circuit.

### Characterization of contralateral heteronymous recurrent inhibition

In the contralateral condition, 59 motor units were recorded, of which 52 exhibited robust inhibition and were therefore retained for subsequent analyses. To investigate contralateral heteronymous RI, we delivered electrical stimulation to the left femoral nerve (contralateral condition, Figure 2A). The stimulation intensity was adjusted to evoke a response equal to 10% of M_max_, targeting motor axonal inputs that may engage polysynaptic commissural pathways, thereby influencing ipsilateral RC activity and modulating right soleus MNs. The analysis revealed a clear heteronymous contralateral recurrent inhibition in the right soleus muscle. The mean latency was 29.9 ± 3.4 ms with an average duration of 44.5 ± 4.2 ms. These results support the presence of a measurable contralateral recurrent inhibition at the motor unit level. When we pooled all motor units recorded across participants and analysed the contralateral condition (Figure 2B), firing rate only moderately explained inhibition duration, similar to the trend observed in the ipsilateral condition (r² = 0.40). Notably, linear extrapolation of inhibition duration to rest predicted a value of 53.6 ms. Individual level analyses using the same predefined r² thresholds used for ipsilateral inhibition revealed a similar, although less pronounced, pattern. We observed strong negative relationships between firing rate and inhibition duration (r² > 0.67) in two participants (S4 and S12). Five participants exhibited moderate relationships (0.33 < r² < 0.67; S1, S3, S5, S7, and S9), while the remaining six participants showed weak or absent relationships (r² < 0.33).

**Figure 2.**
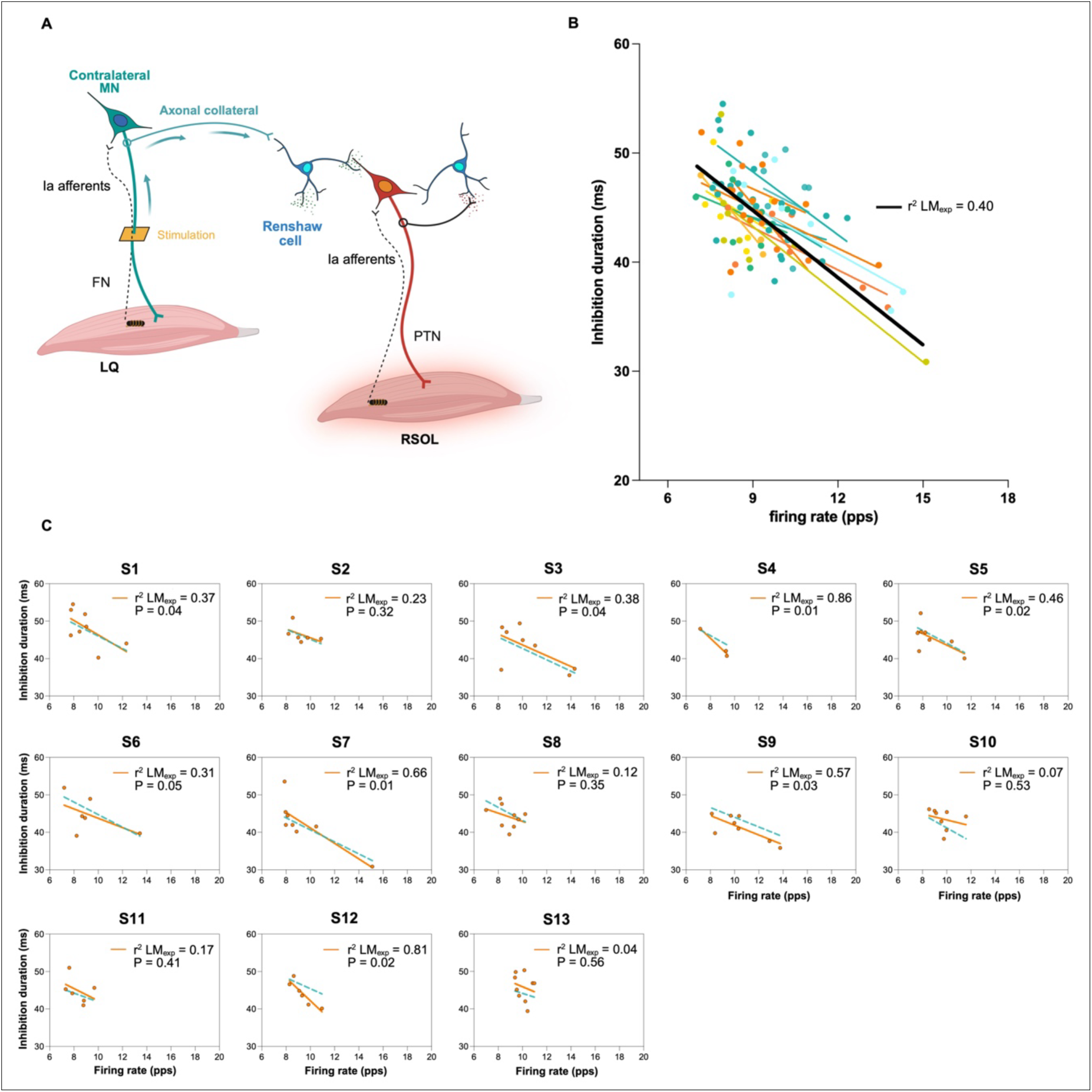
Characterisation of contralateral recurrent inhibition. A) Schematic representation of the contralateral recurrent inhibition pathway (cyan). Electrical stimulation of left quadriceps motor axons was delivered at a constant intensity (approximately 10% of the maximal M-wave) and at intervals between 1 and 2 s, activating commissural MN recurrent collaterals. These collaterals synapse onto Renshaw cells within the spinal cord, which in turn project back onto MNs innervating the right soleus muscle. B) Estimation of contralateral recurrent inhibition duration of single MUs and their firing rate across participants. Colored dots represent MUs and the linear regression fit per individual; black line represents the linear regression fit for the group data with respective r² shown. C) Inhibition duration of individual motor units (MUs) as a function of firing rate for each participant (orange dots). Orange lines represent individualized linear fits (LM_exp_) with the corresponding R² and P values, whereas cyan dotted lines represent the linear mixed-effects model (LMM) regression for each participant. MN, motoneuron; FN, femoral nerve; PTN, posterior tibial nerve, LQ, left quadriceps; RSOL, right soleus.

Regarding inhibition latency, the pooled analysis revealed no relationship between motor unit firing rate and inhibition latency in the contralateral condition (r² = 0.03). Consistently, no participant exhibited a significant relationship at the individual level. Across all individuals, the coefficient of determination remained below the predefined threshold (all r² < 0.19), indicating that firing rate did not predict variations in inhibition latency. These results demonstrate that, unlike inhibition duration, the latency of contralateral heteronymous recurrent inhibition remains independent of motor unit firing rate, as observed for the ipsilateral inhibition. Together, the results from both ipsilateral and contralateral conditions indicate a marked interindividual variability in the dependence of inhibition duration on firing rate. This pronounced heterogeneity underscores the limitations of pooled or single subject analyses for characterizing population level effects.

### Comparison of ipsilateral and contralateral recurrent inhibition properties

Figure 3A illustrates the experimental design used to assess heteronymous RI on the ipsilateral and contralateral sides. We identified an average of 4 to 5 motor units per participant across different regions of the soleus muscle in both ipsilateral and contralateral conditions. After identifying RI in both sides, we sought to determine how the properties of this circuit differ between the ipsilateral and contralateral sides.

**Figure 3.**
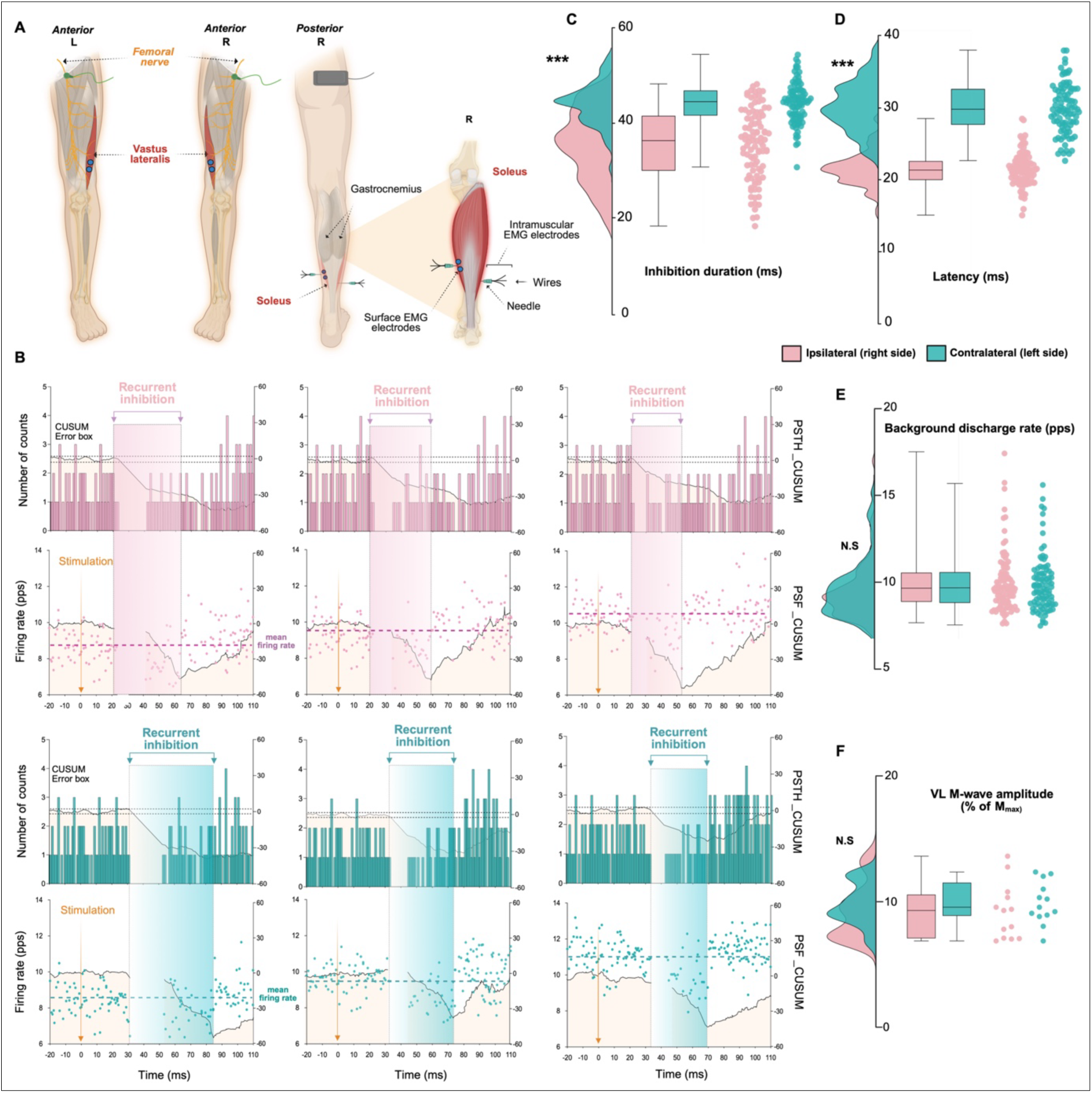
Differential modulation of heteronymous recurrent inhibition in ipsilateral and contralateral conditions. A) Experimental pipeline for assessing ipsilateral and contralateral heteronymous recurrent inhibition in the right soleus muscle. A three-wire intramuscular electrode configuration was used at each implantation site, enabling multiple bipolar recording combinations and expanding motor unit sampling compared with conventional bipolar recordings. To control stimulation parameters, surface electrodes were used to monitor m-wave evoked by stimulation. B) Examples of peristimulus time histograms shown in the top panels and peristimulus frequencygrams shown in the bottom panels from stimulated motor units discharging at different firing rates. Black traces represent the cumulative sum used to quantify deviations from baseline activity. The error box defined by the two horizontal dotted black lines corresponds to the confidence limits used to detect significant inhibitory effects. Shaded regions indicate the time window of recurrent inhibition following stimulation, indicated by the arrow. Dotted horizontal lines represent the mean motor unit firing rate in the ipsilateral condition in pink and in the contralateral condition in cyan. Each column illustrates a different motor unit firing rate, highlighting the frequency dependent modulation of inhibition duration. C) Raincloud plots show the distribution of recurrent inhibition duration for ipsilateral and contralateral stimulation conditions. Each dot represents an individual motor unit, and boxplots indicate the median and interquartile rang. D) Raincloud plots show the distribution of recurrent inhibition latency for ipsilateral and contralateral stimulation conditions. Each dot represents an individual motor unit, and boxplots indicate the median and interquartile range. E) Raincloud plots show the distribution of background motor unit discharge rate for ipsilateral and contralateral stimulation conditions. Each dot represents an individual motor unit, and boxplots indicate the median and interquartile range. F) Raincloud plots show the distribution of the evoked M-wave amplitude expressed relative to M_max_ for ipsilateral and contralateral stimulation conditions. Each dot represents an individual motor unit, and boxplots indicate the median and interquartile range. VL, vastus lateralis; M_max,_ maximal M-wave. ***, P < 0.001; N.S, P > 0.05.

Regarding inhibition latency, LMM analyses revealed a significant main effect of stimulation side. Recurrent inhibition latency was on average 8.5 ms longer in the contralateral than in the ipsilateral condition (F(_1,136_) = 432.98, 95% CI [7.66, 9.26], P < 0.001, Figure 3B and 3D). We did not observe a significant main effect of motor unit firing rate on inhibition latency at the population level (β = 0.014 ms/pps, P = 0.919). Similarly, the interaction between stimulation side and firing rate was not significant (β = –0.30 ms/pps, 95% CI [-0.79, 0.19], P = 0.227), indicating that firing rate did not modulate inhibition latency in either condition. Variance decomposition further revealed that latency variability was driven by differences between individual motor units (∼10%), while variability between participants was negligible after accounting for motor unit–level variability and fixed effects. Although a 10% variance contribution may appear modest, it represents a predominant share of the structured, predictable variance in trial-by-trial spike analyses, where the vast majority of fluctuations are typically driven by residual, stochastic trial-to-trial noise. Within this structured variance, the prominent role of motor unit identity contrasted with the absence of inter-individual effects indicates that inhibition latency is shaped by unique anatomical and structural profiles at the single-unit level. This motor unit-specific signature likely reflects fixed physical features, such as individual axonal conduction distances, peripheral fibre diameters, and localized oligosynaptic pathway configurations, which naturally vary across different units but remain perfectly stable within the same unit across trials.

LMM analyses revealed a significant main effect of stimulation side on RI duration. Inhibition lasted on average 8.7 ms longer in the contralateral than in the ipsilateral condition (F (_1,133_) = 191.81, 95% CI [7.42, 9.88], P < 0.001, Figure 3B and 3C). Using the ipsilateral condition as the reference, we observed a significant effect of firing rate on inhibition duration, with a mean decrease of 2.1 ms per 1 pps increase in firing rate (β = −2.11 ms/pps, P < 0.001). A significant interaction between stimulation side and firing rate (β = 0.996 ms/pps, 95% CI [0.24, 1.75], P = 0.010) indicated that this relationship was attenuated in the contralateral condition, with the inhibition duration decreased by −1.1 ms/pps. Thus, although inhibition duration decreased with firing rate in both conditions, this decrease was significantly less pronounced in the contralateral pathway. Variance decomposition further revealed that both interindividual differences (13.7%) and variability across motor units (11.0%) contributed substantially to total variance.

To verify that differences between ipsi– and contralateral were not driven by differences in motor unit discharge or stimulation parameters, we conducted additional methodological analyses. A paired t-test showed that mean motor unit firing rate did not differ significantly between ipsilateral and contralateral stimulation conditions (P > 0.05, Figure 3E). Similarly, M-wave evoked by stimulation of the femoral nerve did not differ between ipsilateral and contralateral sides (P > 0.05, Figure 3F), as we maintained an evoked amplitude at approximately 10% of maximal M-wave.

Finally, we sought to determine whether the observed RI was modulated by the level of voluntary contraction independently of motor unit discharge rate. Because firing rate increases with force production and is known to influence inhibition duration, a multiple linear regression model was constructed with inhibition duration as the dependent variable and force output, along with the between– and within-motor unit components of firing rate, as predictors. The between-motor unit component reflected differences in average discharge rate across motor units, whereas the within-motor unit component captured deviations from each motor unit’s mean discharge rate across force levels. The resulting model shows a clear negative correlation (R² = 0.463). After accounting for both sources of firing rate variability, force remained a significant predictor of inhibition duration (β = −0.998, P < 0.001), demonstrating that inhibition duration progressively shortened as voluntary contraction intensity increased. These findings indicate that the force-dependent modulation of RI cannot be attributed solely to changes in motor unit discharge rate and likely reflects additional mechanisms associated with increasing motoneuronal drive. This interpretation is consistent with previous reports describing a progressive reduction in RI with increasing levels of voluntary contraction (*9*, *23*).

### In silico modelling of ipsilateral recurrent inhibition properties

One aim of this study was to characterize the properties of heteronymous RI in humans. However, intrinsic limitations of iEMG recordings precluded direct investigation of several key questions, including the neural pathways potentially underlying contralateral RI and the contribution of MN properties to this inhibition. To overcome these constraints, we developed a computational model designed to reproduce our experimental observations and provide a framework for inferring properties inaccessible via non-invasive approaches.

We first implemented a model of ipsilateral RI reproducing the established homonymous Renshaw mediated circuitry (Figure 4A). We modelled all neuronal populations as conductance based leaky integrate and fire units in the Brian2 simulator (*24*). The model simulated a pool of 100 soleus MNs receiving a synaptic drive to reproduce a steady isometric contraction. We modelled MNs to reproduce the electrophysiological properties of soleus motor units, characterized by slow-contracting muscle fibres, including size heterogeneity by assigning soma diameters scaled quadratically from 75 to 85 μm (*25*, *26*). To investigate the mechanisms underlying ipsilateral inhibitory effects we delivered brief phasic activations every 1.8 s to the ipsilateral quadriceps MN pool during the steady state phase of the simulated contraction. To account for interindividual variability, we generated a library of 48 simulations by testing combinations of recurrent inhibitory decay constants (τ_Ri_ = 6, 8, 10, and 12 ms), Renshaw excitation strengths (ω_Ren_ = 0.10, 0.12, 0.14, and 0.16), and quadriceps to Renshaw connectivity probabilities around 17% (14%, 16%, and 18%) (*27*, *28*). For each of these 48 independent library simulations, we randomly selected 20 individual motor units discharging between 6 Hz and 25 Hz for analysis. We processed these simulated data using the exact same PSTH and PSF-based pipeline applied to our experimental iEMG data to ensure direct comparability between in silico and in vivo results. Next, to identify which of the 48 library runs best reproduced the experimental data for each of the 13 participants, we performed a cross-matching procedure. This consisted of comparing the experimental data from each participant against the unit-level relationships of all 48 simulations, selecting the specific simulation run that yielded the lowest mean squared error (MSE) as the participant-specific best-fitting model. To illustrate the comparison between the experimental (LM_exp_) and simulated (LM_sim_) fits (Figure 4B), we first quantified predictive performance using a pseudo r² metric (G), defined as the reduction in prediction error relative to a subject specific null model. For all participants, the best fitting simulation derived models yielded positive G values (Figure 4C), indicating better predictive performance than the subject-specific null model. We further characterized the agreement by comparing the distributions of squared residuals and quantifying differences using Hedges’ g (Figure 4C). In all participants values of g were negligible (g < 0.15) or small (0.15 < g < 0.50), indicating that residual distributions from the simulation derived models closely overlapped with experimental fits. This demonstrates that the model reproduced both the magnitude and the variability of inhibition duration across the recorded range of firing rates. These results indicate that our computational framework, through participant specific selection, robustly captures the individual dependency of ipsilateral RI.

**Figure 4.**
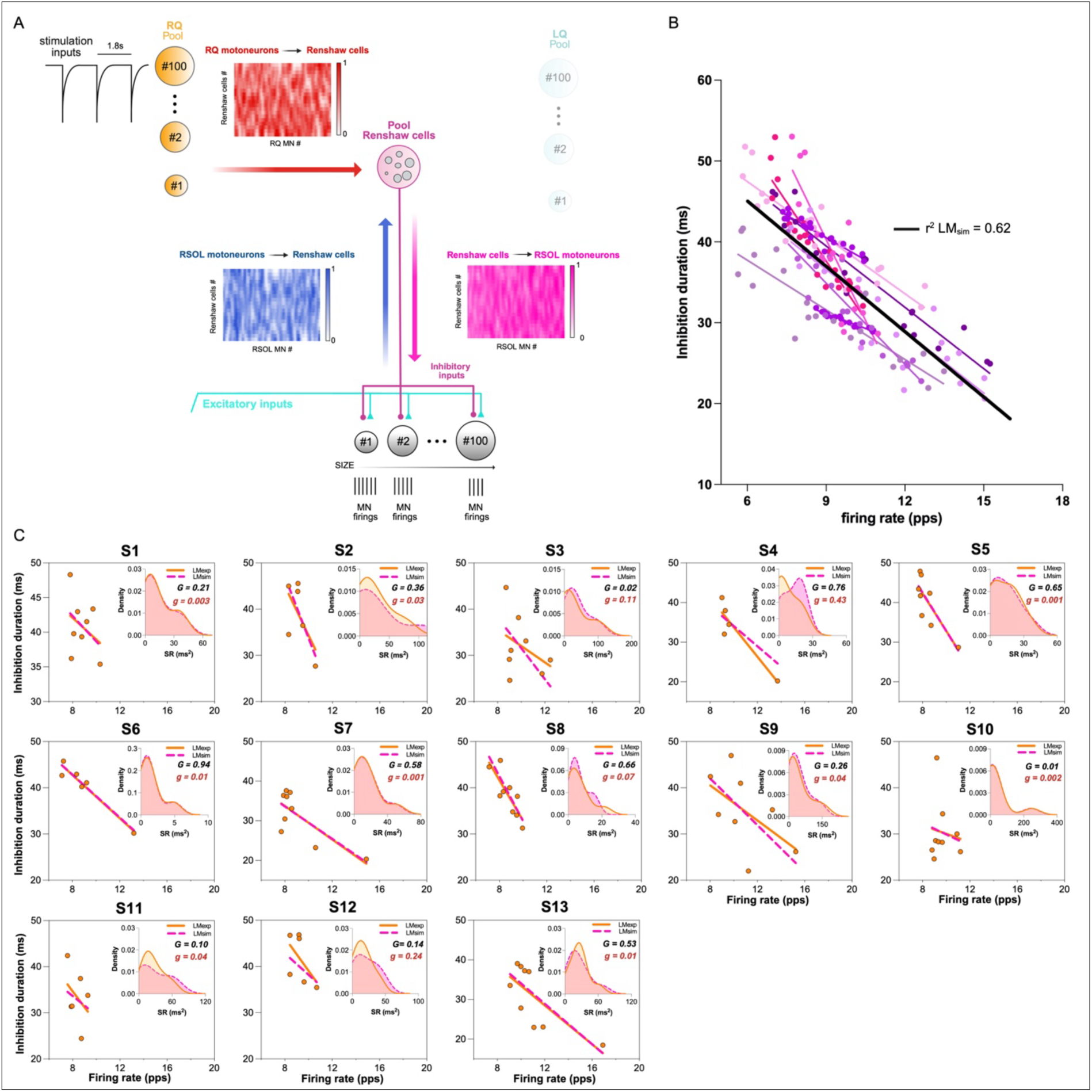
Characterization of ipsilateral recurrent inhibition on individual motoneurons with computational modelling. A) Schematic representation of the in silico computational model illustrating the excitatory drive and the inhibitory input delivered to the right soleus MNs of varying size. The model was used to simulate experimental motor unit firing during isometric voluntary contraction, with periodic inhibitory input originating from quadriceps MNs and delivered to Renshaw cells affecting soleus MNs during the steady state plateau phase of the contraction. Weighted synaptic connectivity matrices illustrating effective connections between motoneural populations, with pixel intensity reflecting normalized cumulative synaptic weight. B) Estimation of inhibition duration and firing rate for individual simulated motor units across the LM_sim_ optimized for each participant. Individual simulated motor units and their corresponding LM_sim_ fits are shown in colour. The black line represents the linear regression fitted to the pooled simulated data, with the corresponding r² value indicated. C) Experimental motor unit firing rate and inhibition duration (orange dots) with the corresponding experimental regression line (orange line; same LM_exp_ as in Fig. 1E) and the participant specific optimal LM_sim_ fit (purple dotted line). Insets show kernel density estimates of the squared residual (SR) distributions for experimental and simulated fits, together with the corresponding pseudo r² (G) and Hedges’ g (g). RQ, right quadriceps. LQ, left quadriceps.

### In silico modelling of contralateral recurrent inhibition properties

To investigate the mechanisms underlying contralateral inhibitory effects, we extended the model to include contralateral stimulation pathways while preserving identical synaptic and intrinsic MN parameters. We delivered brief phasic activations every 1.8 s to the contralateral quadriceps MN pool during the steady state phase of the simulated contraction. We implemented two alternative contralateral network architectures (Figure 5A). In the first one, contralateral quadriceps MNs projected directly to ipsilateral Renshaw cells. In the second, contralateral effects were mediated through a polysynaptic commissural pathway involving V3 interneurons, consistent with anatomical evidence regarding bilateral spinal coordination (*10*). We modelled the intrinsic electrophysiological properties of these V3 interneurons based on the ventral subpopulation identified in in vitro spinal cord slice preparations (*29*). We assigned low leak conductance (0.84 to 1.92 nS) and membrane capacitances (21 to 39 pF) to reproduce the high input resistance and sustained firing patterns characteristic of this ventral cluster.

**Figure 5.**
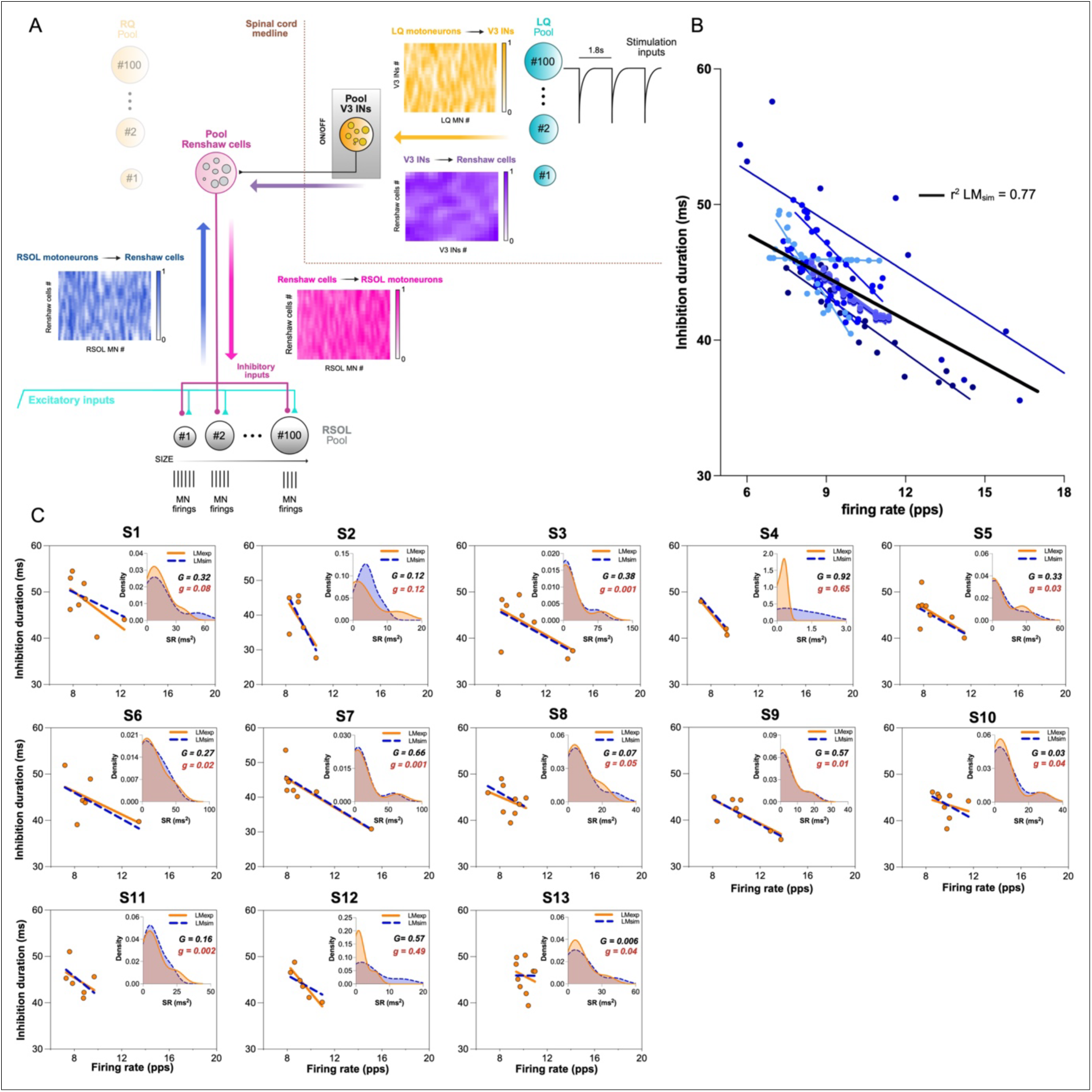
Characterization of contralateral recurrent inhibition on individual motoneurons with computational modelling. A) Schematic representation of the in silico computational model illustrating the excitatory drive and the inhibitory input delivered to right soleus MNs of varying size. The model was used to simulate motor unit firing during isometric voluntary contraction, with periodic inhibitory input originating from left quadriceps MNs and transmitted to Renshaw cells influencing contralateral (right) soleus MNs during the steady state plateau phase of the contraction. In the contralateral configuration, this inhibitory pathway could be implemented either through direct heteronymous projections or via an optional polysynaptic commissural pathway involving V3 interneurons. Weighted synaptic connectivity matrices illustrating effective connections between motoneuronal populations, with pixel intensity reflecting normalized cumulative synaptic weight. B) Estimation of inhibition duration and firing rate for individual simulated motor units across the LM_sim_ optimized for each participant. Individual simulated motor units and their corresponding LM_sim_ fits are shown in colour. The black line represents the linear regression fitted to the pooled simulated data, with the corresponding r² value indicated. C) Experimental motor unit firing rate and inhibition duration (orange dots) with the corresponding experimental regression line (orange line; same LM_exp_ as in Fig. 1E) and the participant specific optimal LM_sim_ fit (blue dotted line). Insets show kernel density estimates of the squared residual (SR) distributions for experimental and simulated fits, together with the corresponding pseudo r² (G) and Hedges’ g (g). RQ, right quadriceps. LQ, left quadriceps.

We generated a library of 96 simulations, comprising 48 runs without commissural V3 interneurons and 48 runs including the V3 mediated polysynaptic pathway. We used the same methodology as for the ipsilateral analyses. Notably, 100% of the best fitting simulation runs across all participants incorporated the V3 mediated pathway (Figure 5B). In twelve participants (S1, S2, S3, S5, S6, S7, S8, S9, S10, S11, S12, and S13), values of g were negligible (g < 0.15) or small (0.15 < g < 0.50), indicating that the simulation derived models closely overlapped with experimental fits (Figure 5C). For the remaining participant (S4), although larger values of g reflected reduced precision in the residual structure, the models still captured the fundamental direction of the firing rate dependency, as evidenced by their positive G values. Together, these results demonstrate that the contralateral RI observed experimentally is best explained by network architectures incorporating commissural V3-like interneurons

### Effect of motoneuron properties on inhibition duration

The duration of inhibition depends on both the strength of inhibitory inputs and the intrinsic properties of MNs that govern synaptic integration. Specifically, MN size critically influences excitability. Larger MNs, characterized by higher membrane capacitance, require greater synaptic currents to produce a given change in membrane potential than smaller MNs. To estimate whether MN properties could influence RI in our experimental dataset, we performed a linear mixed-effects model using motor unit recruitment threshold as a proxy for motoneuron size (*13*). The analysis revealed a significant main effect of recruitment threshold on inhibition duration (F (_1,55_) = 142.47, P < 0.001), with higher-threshold motor units exhibiting shorter inhibition durations (β = −1.05 ms per recruitment threshold unit, 95% CI [−1.23, −0.88]). Importantly, the interaction between recruitment threshold and stimulation side did not reach significance (P = 0.110), suggesting that the association between recruitment threshold and inhibition duration was preserved across sides. Although recruitment threshold is commonly used as an indirect proxy for motoneuron size, it does not provide a direct estimate of the intrinsic biophysical properties.

To address this limitation, we next used model-based biophysical inference in a population of in-silico MNs, allowing direct manipulation and quantification of intrinsic membrane properties, including soma capacitance and its associated conductance-related parameters (Figure 6A).

**Figure 6.**
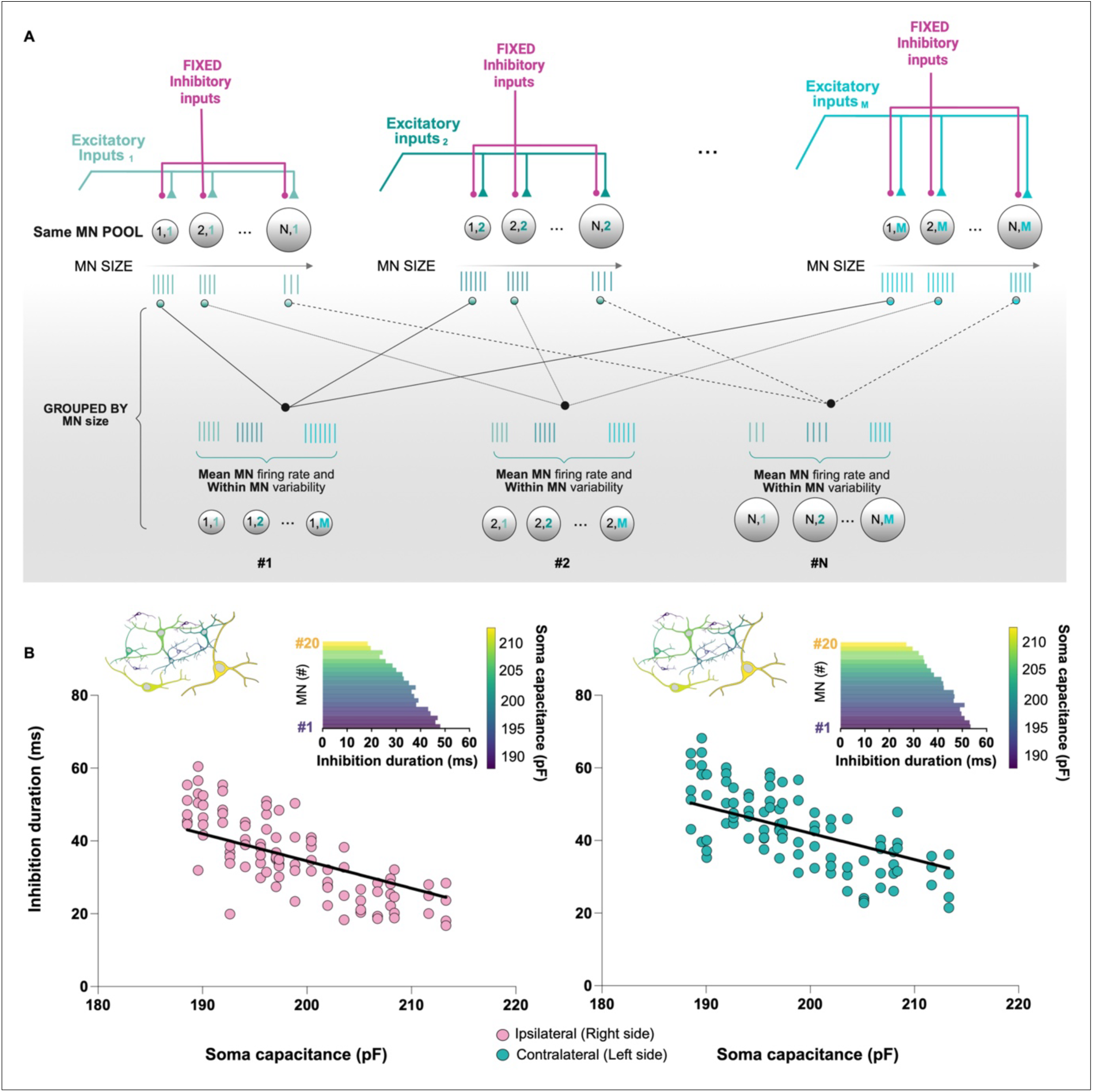
Motoneuron soma capacitance modulated inhibition duration. A) Schematic illustrating our model-based inference approach used to estimate size dependent effects on recurrent inhibition duration. Excitatory input was varied to generate different motor unit firing rates for each MN across a range of sizes, under fixed synaptic parameters and with firing rate decomposed into a unit-specific mean firing rate and a within-unit component. B) Relationship between soma capacitance and recurrent inhibition duration estimated using model-based inference with parameters derived from participant 7, for ipsilateral (pink dots) and contralateral (cyan dots) sides. Solid lines indicate predictions from the linear mixed-effects model, which controls for unit-specific mean firing rate and within-unit firing rate components. Insets illustrate the distribution of inferred inhibition durations across MNs according to soma capacitance.

To examine how intrinsic MN properties shape inhibition duration, we simulated 20 MNs using the in-silico models described previously. We conducted simulations at five different excitatory input levels (corresponding to 2, 5, 10, 15, and 20% MVC) using fixed inhibitory synaptic parameters derived from a representative participant for both the ipsilateral side (τ_Ri_ = 8 ms, ω_ren_ = 0.14, connectivity = 0.16) and the contralateral side (τ_Ri_ = 8 ms, ω_ren_ = 0.14, connectivity = 0.16 with commissural V3 interneuron projections). We tracked individual motor units across increasing contraction levels, spanning a wide range of discharge frequencies for each unit under identical synaptic conditions.

To further examine whether the relationship between RI and MN properties was influenced by firing rate, we fitted a linear mixed-effects model with inhibition duration as the dependent variable. Soma capacitance was included as a fixed effect indexing MN size, whereas side (ipsilateral vs. contralateral) was included to assess potential differences between recurrent pathways. To account for the influence of firing rate, it was decomposed, as in the previous analysis, into between– and within-motoneuron components and included as a covariate in the model. MN identity was included as a random intercept to account for repeated observations within individual units.

The model revealed a robust negative effect of soma capacitance on inhibition duration, indicating that larger MNs exhibited shorter inhibition durations (β = −1.19 ms/pF, 95% CI [−1.49, −0.88], P < 0.001). This effect remained significant after accounting for both firing rate components. Within-MN firing rate deviations were strongly associated with inhibition duration (β = −0.85 ms/pps, P < 0.001), whereas mean firing rate was not (P = 0.930). No significant interactions were observed between soma capacitance and either mean firing rate (P = 0.241) or within-MN firing rate (P = 0.091), indicating a consistent size-dependent effect across firing rate levels.

We also observed a significant effect of laterality (Figure 6B), with shorter inhibition durations on the ipsilateral side compared with the contralateral side (β = –6.77 ms, P < 0.001). However, the interaction between soma capacitance and side was not significant (p = 0.158), indicating that the difference in inhibition duration between ipsilateral and contralateral conditions was not explained by MN size. The mixed-effects model explained a substantial proportion of the variance in inhibition duration (marginal r² = 0.74; conditional r² = 0.84), with approximately 38% of the total variance reflecting baseline heterogeneity across MNs (ICC = 0.38). Together, these results demonstrate that MNs properties, like soma capacitance, contribute to inhibition duration beyond firing rate effects.

## Discussion

The present findings indicate that heteronymous RI in humans extends beyond the classical ipsilateral pathway to include a distinct contralateral component. Intramuscular recordings identified robust inhibitory responses in soleus MNs after ipsilateral and contralateral quadriceps stimulation, with the contralateral pathway characterized by systematically longer latency and duration. Inhibition duration decreased with increasing firing rate in both ipsi– and contralateral stimulation. Moreover, inhibition duration exhibited a marked force-dependent modulation, becoming progressively shorter with increasing levels of voluntary contraction, that likely reflect, in part, the dependence of RI on the MU recruitment threshold. Computational modelling further suggests that the observed contralateral recurrent inhibition is most consistently explained by a polysynaptic commissural pathway involving ventral V3-like interneurons. Furthermore, intrinsic MN properties, particularly soma capacitance and its linked passive parameters, might shape inhibition duration. Together, these observations support the existence of a functional commissural recurrent inhibitory circuit that may contribute to bilateral motor coordination in humans.

### Ipsi– and contralateral heteronymous RI

The ipsilateral RI is characterized by a latency of ∼22 ms and a duration of ∼36 ms for active motor units. This duration is similar to those reported in humans with the M-only method by Özyurt et al. (*13*) during voluntary contraction of the soleus muscle (∼32 ms). When extrapolated at rest to overcome the effect of discharge rate on RI duration, the mean duration is ∼47 ms, close to the value (∼45 ms) obtained by Topkara Arslan et al. (*30*). These durations are also consistent with those reported in other animal models (*31*). It is worth noting that compared with previous work in humans, which primarily focused on homonymous pathways, the present study investigated a heteronymous pathway of RI. The fact that this heteronymous pathway exhibits a duration comparable to those previously established for homonymous responses strongly supports a common RI origin for the transient decrease in discharge probability following motor axon stimulation. Our observations regarding the negative correlation between contraction intensity and inhibition duration provide additional support for RI, as this relationship corresponds to one of its distinctive measurable functional signatures (*9*, *23*). The longer latency of the contralateral heteronymous RI compared with the ipsilateral one (+8.4 ms) was expected due to the longer path and the possible greater number of spinal interneurons mediating the signal transmission from one side to the other (*15*, *32*). For instance, Beith (*32*), while investigating homonymous and heteronymous reflex connections of the paraspinal muscles, reported an increased latency of 9 ms for the contralateral compared with the ipsilateral response. The authors suggested that this longer latency reflected the involvement of additional interneurons. Furthermore, we observed a longer inhibition duration (+8.7 ms) for contralateral RI, which was less expected. This prolonged inhibition may reflect the presence of a polysynaptic commissural pathway potentially contributing to a prolonged temporal profile of inhibition (*10*).

### Dependency of RI on discharge rates

Interestingly, we report a dependency of inhibition duration on MU discharge rates for both the ipsilateral and contralateral heteronymous RI, similar as those reported for homonymous RI (*13*) in both alpha and gamma MN (*31*). This should reflect a general feature of neuronal functioning. The excitability of a neurone is shaped not only by its membrane potential but also by the timing of changes in synaptic conductance (*31*). An inhibitory input produces a change in conductance that outlasts the peak of the inhibitory postsynaptic potential (IPSP) it generates. If a concurrent excitatory input triggers an action potential, the resulting membrane reset reduces the influence of the ongoing IPSP. In other words, the following action potential reduces the influence of the remaining IPSP on subsequent motoneuron discharge (*33*). Furthermore, previous work has shown that the effect of RI depends on its timing within the interspike interval, with stronger effects when inhibition occurs late in the interval, particularly at lower firing rates (*12*). Consequently, when excitatory drive increases, the associated increase in the discharge rate should reduce the duration of inhibition. Importantly, the interindividual and inter-unit variability of the relation between RI duration and firing rate observed in our data, although salient, appeared relatively limited. This is consistent with previous work showing that structured inhibitory circuits display consistent behaviour and weaker dependence on participant specific factors (*21*).

### Motoneuron properties influence RI

RCs generate large inhibitory conductance capable of strongly attenuating MN firing, with a single RC action potential sufficient to transiently silence a MN (*1*). RI thus represents an effective mechanism for modulating MN excitability, shaping firing patterns, and stabilizing motor output. However, a key mechanistic consideration in interpreting inhibitory responses concerns the influence of intrinsic MN properties.

Our in-silico model indicates that membrane capacitance significantly affects the duration of heteronymous RI, independently of the firing rate. Previous studies reported that RI-related IPSP amplitude decreases from low-to high-threshold MNs at rest but becomes similar near firing threshold (*34*), and that differences across MN types may diminish under comparable discharge conditions (*31*). More broadly, intracellular recordings have shown that IPSP amplitude is not directly determined by MN size, suggesting that inhibitory strength primarily reflects synaptic circuit organization (*35*). However, by combining single motor unit recordings with biophysically grounded modelling, we show that inhibition duration depends strongly on intrinsic MN dynamics. Critically, synaptic connectivity and inhibitory input were held constant in the model, isolating intrinsic contributions. Within this framework, membrane capacitance governs the instantaneous rate of membrane potential change at the exact onset of the stimulation, such that a higher capacitance initially buffers the hyperpolarizing impact in larger MNs. However, the subsequent rate of recovery following this transient hyperpolarization and the progression back toward the firing threshold are dictated by the leak conductance and the passive membrane time constant. Because leak conductance scales supralinearly with the square of the soma diameter while capacitance increases only linearly with the MN size, larger motoneurons possess a shorter, faster membrane time constant. Accordingly, the lower capacitance amplifies the initial transient impact of inhibition during the interspike interval in smaller MNs while the combination of a larger capacitance and a faster time constant in larger MNs shortens the inhibition in larger MNs. However, passive membrane properties alone do not fully determine the recovery trajectory between spikes, as this interaction is further shaped by spike-frequency adaptation. In addition to these parameters, other intrinsic properties may also contribute to the membrane trajectory between discharges, particularly the afterhyperpolarization (AHP). Implemented as an activity-dependent current, the AHP defines the membrane trajectory following each spike, such that the effect of inhibition depends on its interaction with this history-dependent state. Stronger AHP (e.g., in small MNs) slows the recovery toward threshold, and thus prolongs the effective impact of inhibition, whereas the faster recovery and shorter time constant in larger MNs reduce the effective impact of inhibition (*36*).

Together, passive membrane properties and AHP kinetics could shape the interspike trajectory and thus determine how inhibitory inputs delay MN spike generation. In this framework, inhibition duration reflects the delay imposed on spike timing, with spike reset limiting the influence of ongoing inhibitory conductance. This is consistent with the negative relation between the duration of the inhibition and the motor unit firing rate observed with experimental data. However, differences in synaptic organization may also contribute to these effects. In particular, low-threshold MN have been reported to receive denser RC projections, both in terms of the number of projecting RCs and the number of synaptic contacts per RC, compared with higher-threshold MN (*35*, *37*).

### V3 commissural interneurons in contralateral heteronymous RI

Previous work suggests that crossed inhibitory pathways in humans may share functional features with commissural networks described in other animal models, particularly those involving V3 interneurons and Renshaw-mediated recurrent circuits (*15*). Our results strongly support the involvement of V3-like commissural interneurons in contralateral heteronymous RI. More specifically, our computational model incorporating a V3-mediated commissural pathway systematically reproduced the main experimental observations across participants and conditions, whereas alternative contralateral models lacking this pathway failed to account for the observed inhibitory dynamics. Consistent with this interpretation, evidence from cross-species locomotor studies highlights the central role of commissural interneurons in shaping interlimb coordination (*19*, *38*). Among these networks, V3 interneurons, which are predominantly glutamatergic, project extensively across the spinal midline and provide excitatory input to contralateral MNs and interneurons, including RC (*19*, *39*, *40*). As glutamatergic neurons, V3 interneurons are expected to recruit both AMPA– and NMDA receptor-mediated transmissions in their target populations. In RC, such inputs could involve slower NMDA-dependent components, potentially contributing to a broader temporal integration of synaptic inputs, and thus influence the duration of contralateral inhibition observed in the present study (Figure 7). In contrast, ipsilateral RI, more directly mediated by MN collaterals onto RC, likely involves a combination of cholinergic and glutamatergic transmission, predominantly engaging fast nicotinic and AMPA receptor-mediated mechanisms (*10*, *13*). While NMDA-dependent currents may contribute to both pathways, the contralateral recruitment of an additional V3-mediated excitatory relay may increase the relative contribution of slower NMDA-mediated components. While the direct involvement of V3 interneurons in RI has not yet been experimentally demonstrated, including in animal models, our results provide the first indirect evidence supporting this possibility. More broadly, comparative neurobiology shows that commissural interneurons are evolutionarily conserved across vertebrates and consistently mediate bilateral coordination of rhythmic behaviour, from fish to humans (*41*). Our work complements and extends existing spinal circuit models by strengthening the case for a commissural inhibitory mechanism underlying contralateral MN modulation through RI, consistent with decades of animal research and emerging human neurophysiological evidence. Furthermore, our study adds new insights to the neural pathway mediating bilateral coordination of rhythmic behaviour.

**Figure 7.**
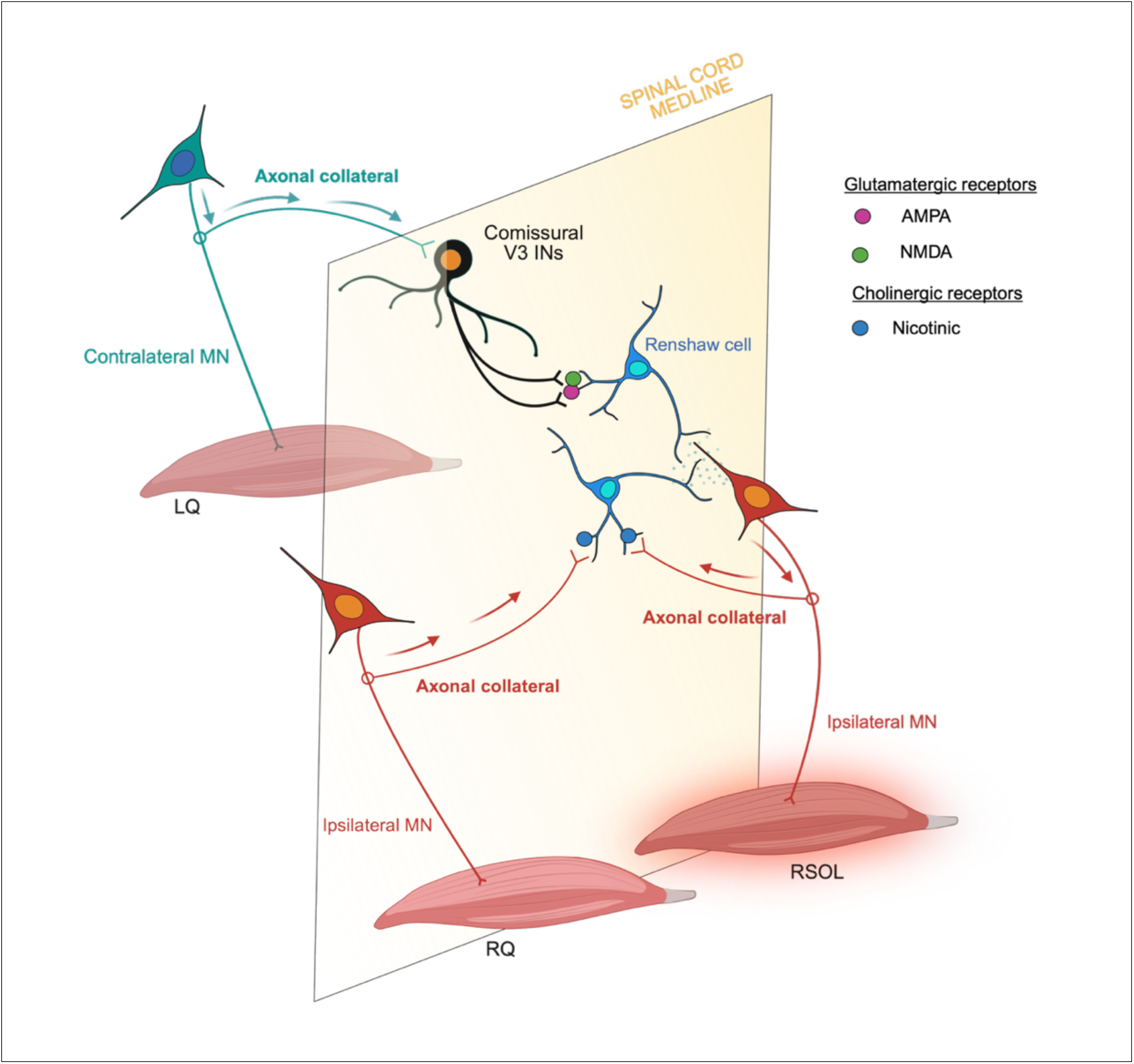
Proposed spinal circuitry underlying ipsilateral and contralateral heteronymous recurrent inhibition. Schematic representation of the spinal circuitry mediating homonymous and heteronymous ipsilateral (in red), as well as contralateral heteronymous (in cyan) recurrent inhibition (RI). Activation of an ipsilateral MN pool in right soleus muscle (RSOL) gives rise to recurrent inhibition via Renshaw cells, which receive excitatory input from MN axon collaterals and provide inhibitory feedback onto the same (homonymous RI) or synergistic (ipsilateral heteronymous RI) MN pools. In addition to these well-described pathways, the present model proposes a commissural mechanism underlying contralateral heteronymous RI. Excitatory commissural V3 interneurons (INs), which project across the spinal cord midline, relay signals originating from motor and/or afferent inputs associated with the activated ipsilateral motor pool to contralateral Renshaw cells. These, in turn, inhibit contralateral MNs, giving rise to a delayed and prolonged inhibitory effect. The contralateral pathway is likely polysynaptic and involves glutamatergic transmission, potentially engaging both AMPA and NMDA receptor-mediated mechanisms, which may contribute to the longer latency and duration of contralateral inhibition. In contrast, ipsilateral RI is more directly mediated and relies more strongly on fast synaptic mechanisms, resulting in a shorter inhibitory response. Arrows indicate excitatory projections, whereas filled circles represent inhibitory synapses. The dashed vertical plane represents the spinal cord midline.

### Methodological considerations

The M-only method captures RC-mediated synaptic input indirectly through decreases in single motor unit discharge rate. Because these inhibitory effects are assessed during voluntary activity, the duration and magnitude of the inhibition reflect not only the underlying RC-induced IPSP but also the state of MN excitability that depends on its discharge rate (*42*). Accordingly, the duration of the decrease in MU discharge rate in response to the stimulation reflects the functional impact of recurrent inhibitory synaptic currents on active MNs, providing a more realistic index of how Renshaw circuitry shapes motor output in vivo (*30*). In addition, previous work by Katz and Pierrot-Deseilligny (*9*) showed that RI may be modulated differently depending on the muscle investigated and the motor task performed. Therefore, although the present study demonstrates a clear relationship between motor output and inhibitory responses in the soleus muscle, these findings may not necessarily generalize to all muscles or behavioural contexts. Despite these limitations, the inhibitory effects evoked by femoral nerve stimulation onto soleus motor units may reflect a more functionally relevant pattern of spinal inhibitory interactions than classical homonymous RI paradigms. Indeed, the present paradigm involves heteronymous and contralateral interactions that are likely to contribute to interlimb coordination during movement (*43*). Furthermore, despite the advantages of the M-only approach, the present study still relies on indirect measures of recurrent inhibitory circuitry and therefore retains inherent methodological limitations. Finally, we cannot exclude that the stimulation, even in the absence of H-reflex, may have concurrently recruited the most excitable afferents, including Ia and potentially Ib afferents. Although group Ib afferents generally require higher activation thresholds (*44*) and should be limited, the resulting responses can reflect a mixture of excitatory and inhibitory postsynaptic influences rather than isolated RI. Taken together, although these methodological considerations warrant cautious interpretation, the convergence of the experimental observations and computational modelling supports recurrent inhibition as the most likely mechanism underlying the inhibitory responses observed in this work.

The present study demonstrates that heteronymous RI in humans includes not only the classical ipsilateral pathway but also a distinct contralateral inhibitory component characterized by longer latency and duration. By combining intramuscular recordings with computational modelling, we show that the temporal profile of RI is shaped jointly by spinal circuit organization and intrinsic motoneuron properties, including passive membrane dynamics and activity-dependent mechanisms. Our findings further support the involvement of a polysynaptic commissural pathway consistent with V3-like interneuronal networks, providing the first indirect evidence that commissural recurrent inhibitory circuits may contribute to bilateral motor coordination in humans. Together, these results extend the current understanding of RI beyond local ipsilateral feedback control and suggest that spinal inhibitory networks participate in the dynamic coordination of bilateral motor output.

## Materials and Methods

### Ethical approval

The described research was in accordance with the latest revision of the Declaration of Helsinki and was approved by the Ethics committee of Erasme Hospital (#P2025/232). Participants provided oral and written consent prior to experimental testing.

### Participants

Thirteen volunteers (four females and nine males), with a mean age of 25 ± 3 years, a height of 175 ± 11 cm, and a body mass of 74 ± 17 kg, participated in this study. Verbal screening was used to exclude individuals with neuromuscular or metabolic disorders. Participants were also instructed to avoid strenuous physical activity for 48 hours prior to each testing session.

### Experimental Design

The objective of this study was to determine the influence of contraction intensity on the ipsilateral recurrent inhibition pathway and to determine whether a contralateral recurrent inhibition pathway may exist in humans. We further sought to consider whether the contralateral pathway involves commissural V3 interneurons. Additionally, because the strength of the inhibition across contraction intensities can be influenced by MN size and discharge rate, we dedicated a silico model to explore these aspects. To achieve this, we combined iEMG recordings with peripheral nerve stimulation during isometric contractions (experimental data) and designed a biophysical in silico model to isolate MN size and discharge rate as potent biases that are not directly accessible in human participants.

We seated participants upright in a chair with their right leg positioned in a custom-built dynamometer designed to measure isometric plantar flexion force. We aligned the hip, knee, and ankle joints at 90°, with a force transducer (Kistler, Type 4576A1NC1) above the knee. During each effort, participants lifted their heel to produce a plantar flexion movement. We converted the resulting signal from analogue to digital using a Power1401 (Cambridge Electronic Design, Cambridge, UK) and sampled it at 1000 Hz using Spike2 software. A monitor placed approximately one meter in front of the participant provided real time visual feedback of the force output.

First, subjects performed three 3-s plantar flexor maximal voluntary isometric contractions (MVC), with 1-min rest intervals. For each trial, the mean force over the 500-ms window corresponding to peak force was calculated, and these values were averaged across trials to define 100% MVC for each participant. Participants then completed a series of triangular isometric contractions designed to characterize motor unit recruitment behaviour. Each trial began at 0% MVCi, increased progressively to 20% MVCi, and then returned to baseline in order to maintain stable and reliable motor unit decomposition while minimizing discomfort and fatigue for the participants. The force ramp increased and decreased linearly over 10s to promote controlled and gradual modulation of force output. This protocol enabled accurate identification of individual motor unit recruitment thresholds by determining the force level at which motor units began discharging during the ascending phase of the contraction.

After determining recruitment thresholds, participants performed a series of prolonged isometric contractions, each lasting ∼5 minutes. These sustained contractions were performed at three to four different force levels selected to isolate specific single motor unit action potentials in real time using Spike2 software combined with an oscilloscope, thereby enabling the assessment of RI in identified motor units. We interspersed these sustained contractions with 5-min rest periods to minimize fatigue effects.

### Electromyographic recordings

Bipolar surface electromyography was recorded from vastus lateralis (VL) muscles using silver chloride electrodes (8 mm diameter), with an interelectrode distance of 20 mm. To minimize impedance at the skin electrode interface, we shaved the skin when necessary and cleaned it with a solution of alcohol, ether, and acetone. Following the SENIAM recommendation, the VL electrodes were fixed lengthwise 2/3 on the line from the anterior superior iliac spine to the lateral side of the patella. We placed reference electrodes on the skin over the right tibia. We amplified the EMG signals (×1000) and band pass filtered them between 10 and 1000 Hz. We sampled the data at 2 kHz using a Power1401 system (16-bit resolution; Cambridge Electronic Design) and stored the signals on a computer for subsequent analysis.

### Intramuscular recordings

We recorded single motor unit potentials from the SOL muscle using stainless steel wires (50 μm diameter) insulated with Formvar (California Fine Wire, Grover Beach, CA, USA). Each electrode included three wires to increase the number of available recording combinations. We inserted the wires into the muscle belly using a 27-gauge hypodermic needle, which we removed once the wires were positioned. For each session, we implanted two to three pairs of electrodes at different locations along the muscle belly to increase the number of motor units (*45*).

We positioned a reference electrode for both surface and intramuscular EMG on the skin over the patella. We amplified the single motor unit potential recordings (×1000) and band pass filtered them between 10 Hz and 5 kHz. We sampled the motor unit signals at 20 kHz using a Power1401 interface (16-bit resolution; Cambridge Electronic Design) and stored them on a computer. We subsequently identified the single motor unit potentials offline using Spike2 software (Cambridge Electronic Design).

### Peripheral nerve stimulation

To evoke heteronymous recurrent inhibition, we used the “M only stimulation” method described by Ozyurt et al. (*13*), which selectively activates the motor axons of the nerve while minimizing sensory fibre recruitment. Percutaneous stimulations were applied on the right and left femoral nerves with a single rectangular pulse delivered by a constant current stimulator (pulse duration: 0.2 ms; Digitimer stimulator, model DS7, Hertfordshire, UK). The cathodes (AG-Cl, 8 mm diameter) were positioned on both femoral triangles, in the location that elicited the maximum quadriceps twitch amplitude at rest. The anodes (5 cm × 10 cm, Medicompex SA, Switzerland) were placed midway between the right great trochanter and the iliac crest. The pulse duration was reduced to 0.2 ms relative to longer durations (e.g. 1 ms) commonly used in M-only paradigms, to bias stimulation toward motor axons over sensory afferents, consistent with their shorter chronaxie and distinct strength–duration properties (*46*). This intermediate duration ensured reliable M-wave responses without requiring high stimulation intensities, thereby limiting discomfort and minimizing unintended afferent recruitment.

We established the M-wave threshold and the maximal M-wave amplitude (M_max_) for both VL muscles at rest. M-wave threshold was defined as the lowest intensity eliciting a detectable M-wave. We standardized the stimulation intensity for the M only protocol to evoke an M-wave amplitude corresponding to approximately 10% of M_max_, ensuring either no H-reflex or the presence of a minimal H-reflex. Conversely, we determined M_max_ by increasing the stimulation intensity until no further increase in M-wave amplitude occurred for three successive intensities.

Once we observed a stable discharge rate for the isolated motor units, we applied M only stimulation to the femoral nerve. We delivered these stimuli in a random sequence with an interstimulus interval between 1 and 2 seconds. Each participant contributed recordings from at least three distinct motor units. For both ipsilateral and contralateral sessions, the total number of stimuli delivered ranged from 500 to 1000.

#### Spiking neural network model

We simulated all neural populations using conductance based leaky integrate and fire models implemented in the Brian2 simulator for Python (*24*). We integrated membrane voltage dynamics using an explicit Euler scheme with a fixed 0.1 ms time step (*47*). The architecture reproduced core components of spinal motor circuitry relevant to our experimental paradigm. This included two quadriceps MN pools (right and left; Qd and Qg), one right Sol MN pool, a population of ipsilateral Renshaw cells (Rs), and, depending on the simulated condition, a pool of commissural V3 interneurons.

We simulated two physiologically inspired configurations. Under ipsilateral stimulation, only the homonymous recurrent inhibitory loop from right quadriceps MNs to Renshaw cells was present. Under contralateral stimulation, we activated ipsilateral Renshaw cells either via a direct heteronymous pathway from left quadriceps MNs or through an indirect polysynaptic pathway mediated by V3 commissural interneurons.

Each simulation lasted 80 s and began with a ramp increase from 0 to a level equivalent to 20% MVC, followed by a steady plateau similar to the experimental conditions. During the steady-state plateau, we computed the mean discharge rate of all 100 MNs in the Sol pool and retained units firing between 5 and 25 Hz. From this physiologically relevant subset, we randomly selected 20 MNs and analysed them using the same pipeline applied to the experimental data. This fixed sample size was used to ensure that the same number of units was analysed across simulations and to remain consistent with previous modelling work (*21*). All code used for model generation, simulation, and analysis is available at: https://github.com/ (after acceptance).

#### Motoneurons model

We modelled all neurons as conductance based leaky integrate and fire (LIF) units with a membrane reset to 0 mV and a spike threshold of 10 mV, consistent with previous spinal modelling work (*47*). Quadriceps MNs followed the standard LIF equation:

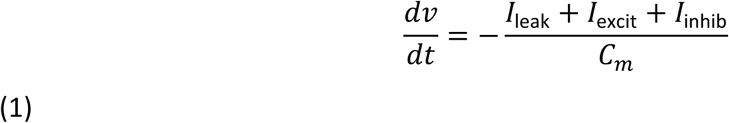

Soleus MNs (i.e., the actively contracting pool) included additional intrinsic mechanisms:

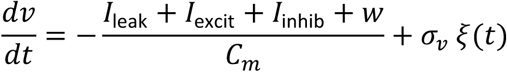

where *w* is the afterhyperpolarization (AHP) current triggered by each spike, and *ξ*(*t*) is Gaussian white noise (amplitude *σ_v_*). This formulation reproduces hallmark electrophysiological features of slow-type motor units, including low recruitment thresholds, relatively regular discharge, spike-frequency adaptation, and modest interspike variability (*25*).

Membrane currents were defined as:

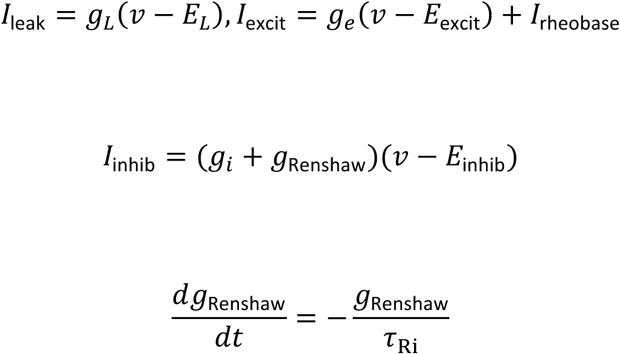

with reversal potentials *E_L_* = 0 mV, *E*_excit_ = +20 mV, and *E*_inhib_ = −20 mV. τ_Ri_ denotes the decay time constant of the Renshaw-mediated inhibitory conductance acting on Sol MNs. MN pools (Qd, Qg, Sol) each contained 100 neurons, whereas Rs and V3 pools contained 20 interneurons each, following the established 1:5 interneuron-to-MN ratio (*47*, *48*). Each MN was assigned a soma diameter:

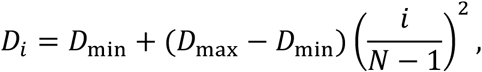

We assigned each MN a soma diameter ranging from a minimum of 75 µm to a maximum of 85 µm. The quadratic exponent used in this distribution yields a physiologically realistic population dominated by small, highly excitable MNs (*26*, *49*). Soma diameter determined the passive membrane properties of the unit. Specifically, membrane capacitance scaled linearly with diameter, whereas leak conductance followed a supralinear relation with the square of the diameter.

This configuration reproduced size dependent recruitment dynamics consistent with experimental observations. After each spike, we reset the membrane voltage to 0 mV for a 10 ms refractory period. We set the reference capacitance to 200 pF, which corresponds to the effective soma capacitance obtained by applying a classical specific membrane capacitance of 1 µF/cm² to a morphologically realistic MN. While Valdunciel et al. (*21*) implemented a two-compartment model, we represented the soma as a single equivalent unit.

#### Synaptic conductance and external drive

Excitatory and inhibitory synaptic conductance evolved dynamically according to:

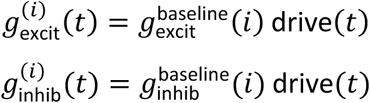

Where (*t*) denotes time, (*i*) denotes the MN, and drive(*t*) denotes the synaptic input applied exclusively to Sol MNs during contraction. The drive comprised: a slowly increasing ramp (5% increments every 2 s up to 20%), low-frequency oscillations (∼1 Hz), and a small stochastic term. This yielded physiologically realistic fluctuations in Sol force output.

Baseline conductance varied across MNs through a size-dependent gradient combined with log-normal multiplicative noise, generating realistic heterogeneity in recruitment thresholds and synaptic gain. Quadriceps MNs did not receive drive and therefore remained at rest unless externally stimulated. Their only activation arose from brief phasic bursts designed to emulate quadriceps stimulation during the Sol plateau. These bursts consisted of high-amplitude injected currents (2 µA, 1-ms duration) repeated every 1.8 s (*21*), applied either to the right or left quadriceps pool depending on the simulated condition. Because quadriceps MNs had no tonic drive, these bursts formed their sole source of depolarization, enabling precise assessment of how recurrent inhibition and commissural pathways transform phasic quadriceps inputs.

#### Muscle fibre model

To confirm that the computational model reproduced physiologically realistic force production, we implemented a muscle fibre model. Each Sol MN innervated a single muscle fibre, which was modelled using standard twitch-like dynamics to reproduce the characteristic rapid rise and slower decay of force observed experimentally in slow muscle fibres. Force production for each fibre followed:

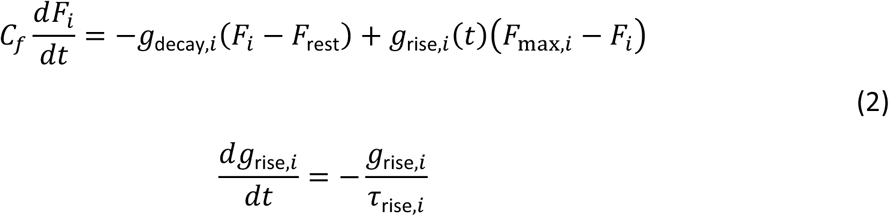

where *g*_rise,*i*_(*t*) represents the activation state of fibre *i*. This variable was incremented by a fixed amount upon each Sol MN spike, thereby reproducing the nonlinear summation and rate-dependent modulation of twitch amplitudes typically observed in slow motor units. The formulation ensures that fibre force displays a sharp onset followed by a slower relaxation, consistent with single-twitch physiology.

Total muscle force was computed as the sum of all fibre forces:

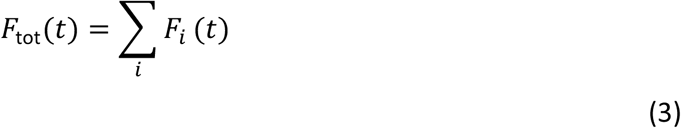

A separate maximal-drive simulation was conducted to determine the model’s maximal force output F_max_, corresponding to full recruitment and maximal firing of the entire Sol MN pool. All plateau forces obtained during the simulated contraction were then normalized to %MVC using:

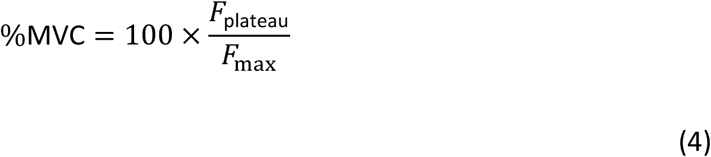

This normalization ensured that force output was expressed in physiologically interpretable units directly comparable to the recorded experimental contractions. The combination of twitch-based fibre dynamics and MVC-based normalization guaranteed that the force produced by the MN pool remained physiologically coherent, enabling the model to generate realistic submaximal contractions at 2–20% MVC. In this way, the muscle model provided a biophysically grounded bridge between MN spiking activity and the targeted force levels imposed in each simulated condition.

#### Recurrent inhibition

Renshaw cells were modelled as:

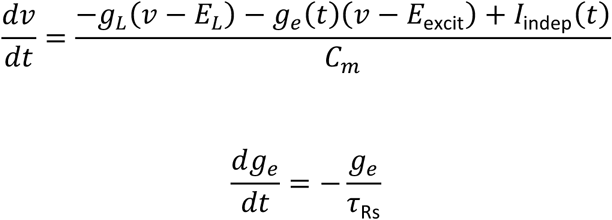

V3 interneurons followed an identical structure but without independent input:

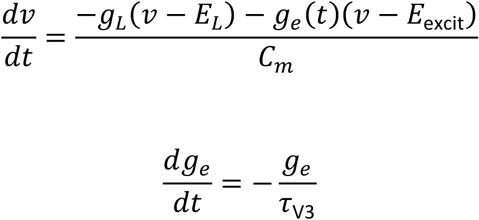

We constrained the passive membrane properties of V3 interneurons, specifically membrane capacitance and input resistance, using electrophysiological recordings from genetically identified V3 interneurons. We selected parameter values to match the most excitable V3 subgroup reported in mouse spinal cord preparations (*29*).

Synaptic decay constants onto Renshaw (τRs) and V3 (τV3) interneurons were fixed at 8 ms (*28*, *50*). Upon reaching 10 mV, cells reset to 0 mV and entered a 10 ms refractory period (*47*). Right and left quadriceps MNs projected to RC with a baseline connection probability of 17% (*28*), around which this parameter was varied in a parameter sweep. To account for homonymous recurrent inhibition, we implemented an additional 17% connectivity from Sol MNs to RC. When V3 interneurons were included, left quadriceps MNs activated RC through a commissural pathway, where 17% of left quadriceps MNs projected to V3 interneurons and 40% of V3 interneurons projected to RC. Each Sol MN received inhibitory input from 40% of the RC population, which is consistent with previous studies (*28*, *47*). The parameter ωren entered the model through the excitatory synaptic update rule onto RC. This scaled the increment of excitatory conductance evoked by each presynaptic spike, allowing for uniform adjustment of recurrent inhibitory strength across the network.

The transmission delay was 5 ms for the right quadriceps projection to RC, 8 ms for the left projection, and 18 ms for the total V3 pathway. Interactions between RC and Sol MNs in both directions used a 5 ms delay (*47*).

#### Biophysical parameters sweep for recurrent inhibition

To characterize how RI duration depends on Renshaw circuit properties, we performed a systematic biophysical parameter sweep. We varied four synaptic decay constants of the Renshaw-to-MN inhibitory conductance (τRi; 6, 8, 10, and 12 ms), four Renshaw excitation strengths (ωren; 0.10, 0.12, 0.14, and 0.16), and three connection probabilities between quadriceps MNs and Renshaw or V3 interneurons (14%, 16%, and 18%, Table 1). For contralateral stimulation only, we also varied the presence or absence of V3 commissural interneurons. We executed all simulations automatically using a Papermill based workflow implemented in Python to ensure reproducibility and comprehensive exploration of the parameter space. This sweep resulted in 48 simulations for the right-side condition and 96 simulations for the left side condition, yielding a total of 144 simulations for the full parameter exploration.

**Table 1.**
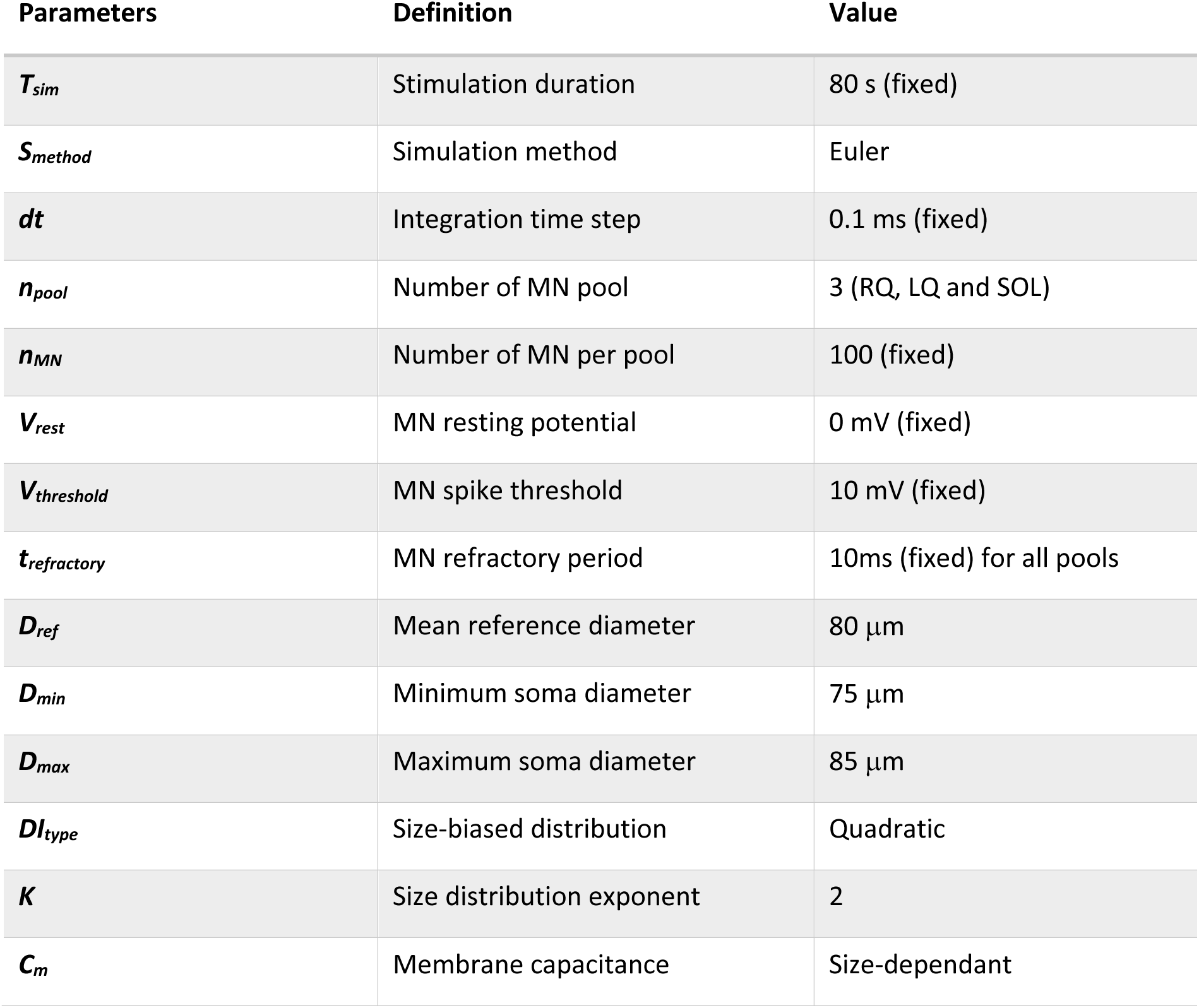
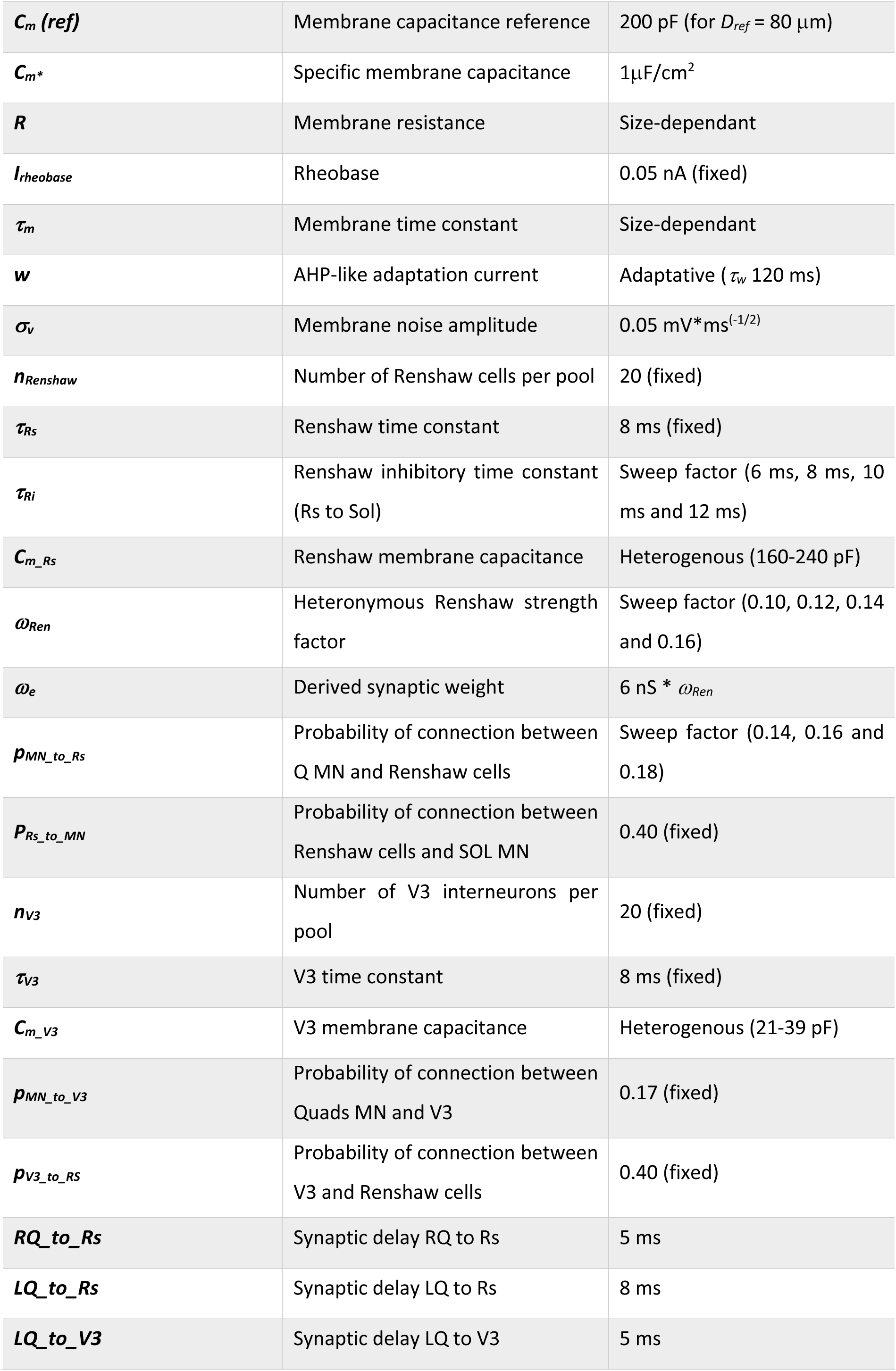
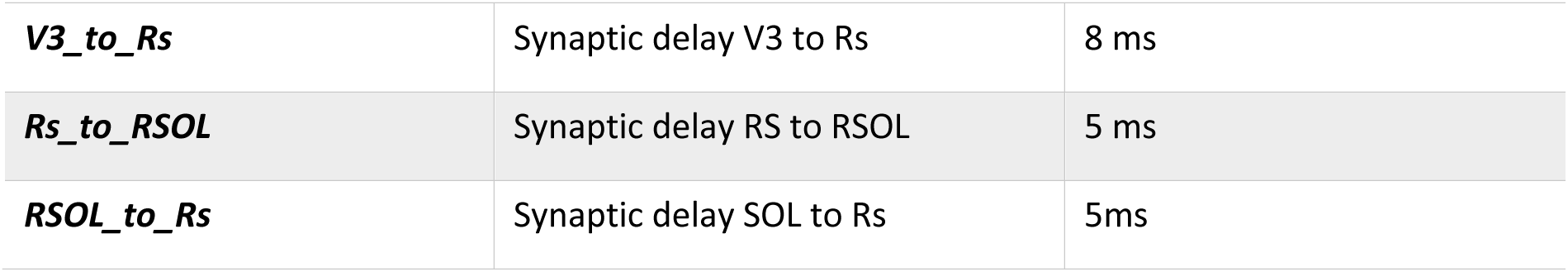
Simulation parameters. RQ, right quadriceps; LQ, left quadriceps; RSOL, right soleus; MN, MNs; Rs, Renshaw cell; V3, V3 interneurons.

| Parameters | Definition | Value |
| --- | --- | --- |
| $T_{sim}$ | Stimulation duration | 80 s (fixed) |
| $S_{method}$ | Simulation method | Euler |
| $dt$ | Integration time step | 0.1 ms (fixed) |
| $n_{pool}$ | Number of MN pool | 3 (RQ, LQ and SOL) |
| $n_{MN}$ | Number of MN per pool | 100 (fixed) |
| $V_{rest}$ | MN resting potential | 0 mV (fixed) |
| $V_{threshold}$ | MN spike threshold | 10 mV (fixed) |
| $t_{refractory}$ | MN refractory period | 10ms (fixed) for all pools |
| $D_{ref}$ | Mean reference diameter | 80 μm |
| $D_{min}$ | Minimum soma diameter | 75 μm |
| $D_{max}$ | Maximum soma diameter | 85 μm |
| $DI_{type}$ | Size-biased distribution | Quadratic |
| $K$ | Size distribution exponent | 2 |
| $C_m$ | Membrane capacitance | Size-dependant |
| $C_m (ref)$ | Membrane capacitance reference | 200 pF (for $D_{ref} = 80 \mu m$) |
| $C_m^*$ | Specific membrane capacitance | $1 \mu F/cm^2$ |
| $R$ | Membrane resistance | Size-dependant |
| $I_{rheobase}$ | Rheobase | 0.05 nA (fixed) |
| $\tau_m$ | Membrane time constant | Size-dependant |
| $w$ | AHP-like adaptation current | Adaptative ($\tau_w$ 120 ms) |
| $\sigma_v$ | Membrane noise amplitude | $0.05 mV \cdot ms^{(-1/2)}$ |
| $n_{Renshaw}$ | Number of Renshaw cells per pool | 20 (fixed) |
| $\tau_{Rs}$ | Renshaw time constant | 8 ms (fixed) |
| $\tau_{Ri}$ | Renshaw inhibitory time constant (Rs to Sol) | Sweep factor (6 ms, 8 ms, 10 ms and 12 ms) |
| $C_{m\_Rs}$ | Renshaw membrane capacitance | Heterogenous (160-240 pF) |
| $\omega_{Ren}$ | Heteronymous Renshaw strength factor | Sweep factor (0.10, 0.12, 0.14 and 0.16) |
| $\omega_e$ | Derived synaptic weight | $6 nS * \omega_{Ren}$ |
| $p_{MN\_to\_Rs}$ | Probability of connection between Q MN and Renshaw cells | Sweep factor (0.14, 0.16 and 0.18) |
| $p_{Rs\_to\_MN}$ | Probability of connection between Renshaw cells and SOL MN | 0.40 (fixed) |
| $n_{V3}$ | Number of V3 interneurons per pool | 20 (fixed) |
| $\tau_{V3}$ | V3 time constant | 8 ms (fixed) |
| $C_{m\_V3}$ | V3 membrane capacitance | Heterogenous (21-39 pF) |
| $p_{MN\_to\_V3}$ | Probability of connection between Quads MN and V3 | 0.17 (fixed) |
| $p_{V3\_to\_Rs}$ | Probability of connection between V3 and Renshaw cells | 0.40 (fixed) |
| $RQ\_to\_Rs$ | Synaptic delay RQ to Rs | 5 ms |
| $LQ\_to\_Rs$ | Synaptic delay LQ to Rs | 8 ms |
| $LQ\_to\_V3$ | Synaptic delay LQ to V3 | 5 ms |
| *V3_to_Rs* | Synaptic delay V3 to Rs | 8 ms |
| *Rs_to_RSOL* | Synaptic delay RS to RSOL | 5 ms |
| *RSOL_to_Rs* | Synaptic delay SOL to Rs | 5ms |

To examine how intrinsic MN properties shape heteronymous RI, we simulated 20 MNs using the biophysical model described above. We conducted simulations at five different excitatory drives (2, 5, 10, 15, and 20% MVC) using fixed inhibitory synaptic parameters derived from a representative participant. For the ipsilateral right side, we set the synaptic decay constant to 8 ms, the Renshaw excitation strength to 0.14, and the connectivity to 0.16. For the contralateral left side, we used identical parameters but included commissural V3 interneuron projections. We tracked individual motor units across increasing contraction levels, spanning a wide range of discharge frequencies for each unit under identical synaptic conditions.

MN size and discharge rate are inherently linked, as size influences recruitment behaviour while discharge rate directly modulates inhibitory efficacy. One strategy to dissociate these factors is to enforce similar firing rates across MNs of different sizes by adjusting excitatory input (*21*). Here, we adopted an alternative approach by preserving the physiological relationship between size and firing rate and accounting for their covariation analytically. Firing rate varied naturally with excitatory drive, and we explicitly controlled its contribution to recurrent inhibition by decomposing firing rate into the MN specific mean firing rate across observations and within MN deviations from this mean. This framework enabled the estimation of size related effects on RI at a constant firing rate without imposing artificial discharge conditions.

### Data Analyses

#### Decomposition

We performed motor unit analysis following established procedures (*51–53*). We initially identified single motor unit action potentials using template matching tools implemented in Spike2 software. An experienced operator visually inspected waveform overlays to verify consistent shape and amplitude within each motor unit train. We generated interval time histograms to confirm physiologically plausible discharge regularity. We only retained units containing at least five consecutive discharges and exhibiting a coefficient of variation of interspike intervals less than or equal to 30% (*54–56*).

When multiple motor units contributed to the recorded signal, we performed an additional decomposition using principal component analysis followed by an ellipsoidal Gaussian mixture model (Figure 8A). This approach probabilistically classified spikes in principal component space and improved sorting accuracy in the presence of overlapping or variable waveforms

**Figure 8.**
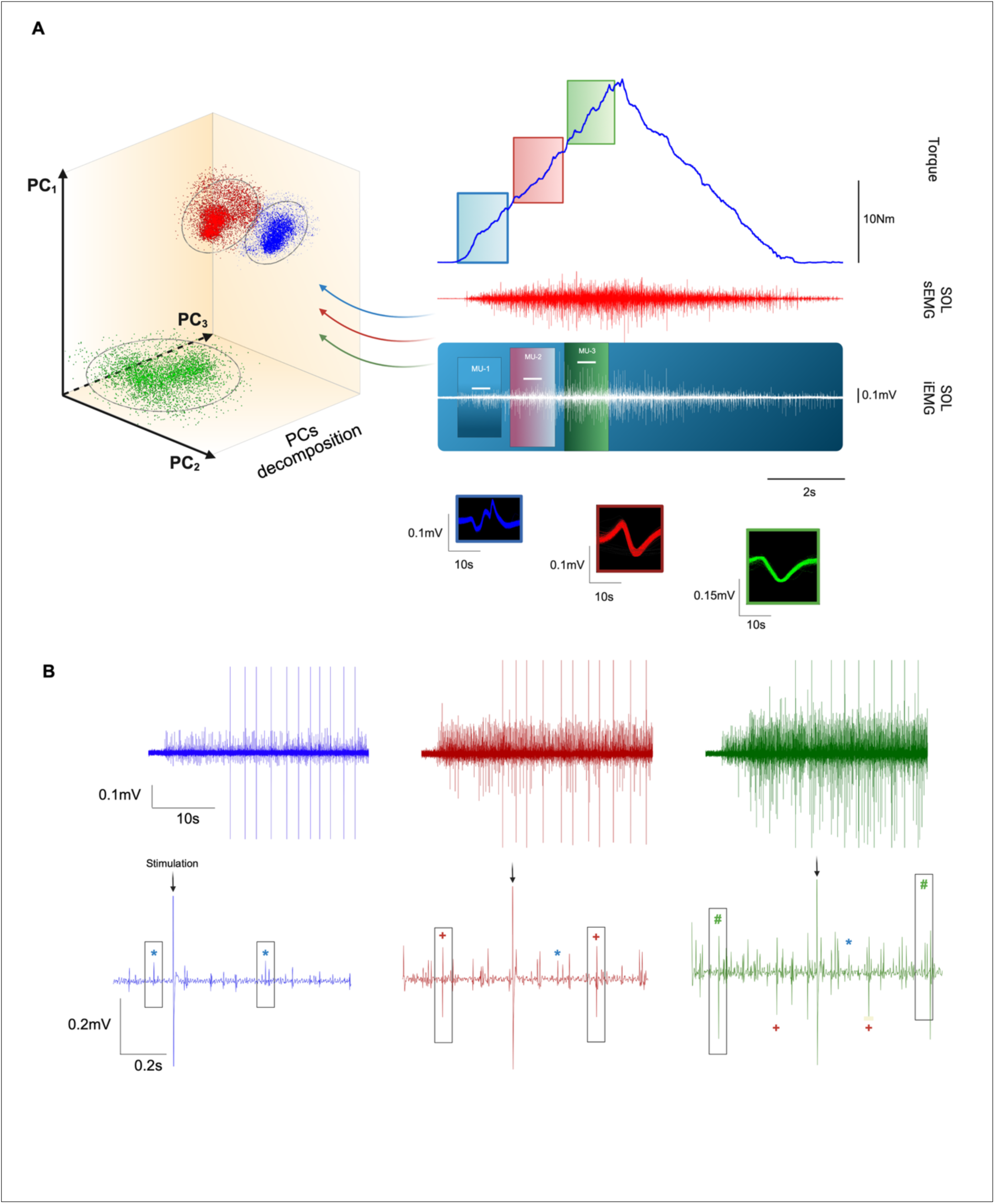
Illustration of the motor unit decomposition and tracking procedure applied during the experiment. A) Identification of motor units and their recruitment thresholds using automated selection algorithms and principal components. B) Tracking of motor unit activity during isometric contractions at different torque levels, with stimulation delivered randomly at 1–2 s intervals. sEMG, surface electromyography; iEMG, intramuscular electromyography; MU, motor unit; PC, principal component; * first motor unit; +, second motor unit; #, third motor unit.

(*57*). We applied manual corrections when necessary to ensure accurate spike assignment. Once isolated, individual motor units were tracked across a range of contraction intensities (Figure 8B).

#### Recurrent inhibition properties

We quantified inhibitory responses from motor unit spike trains using complementary peristimulus time histograms (PSTH) and peristimulus frequency (PSF) analyses (*13*). Briefly, PSTHs were used to estimate inhibition latency (onset), whereas PSFs were used to determine inhibition offset. Both PSTH and PSF were computed with 1 ms bin widths, and cumulative sum (CUSUM) transforms were applied to enhance detection of stimulus-locked deviations from baseline activity.

These metrics provide complementary sensitivity to inhibitory dynamics. PSTH reflects stimulus-related changes in firing probability and is well suited to detect inhibition onset but is less sensitive to sustained inhibition when firing probability recovers despite a reduced discharge rate. PSF tracks instantaneous discharge rate and better captures prolonged inhibitory effects. Combining PSTH-derived onset with PSF-derived offset provides a more accurate estimate of total inhibition duration (*58*). The significance of poststimulus inhibitory events was determined using the CUSUM error box method (*59*).

We estimated baseline variability from the prestimulus interval (−250 ms to 0 ms) and used the largest deviation observed during this period to define symmetrical upper and lower thresholds around the CUSUM trace. These thresholds were used as the criterion for detecting significant synaptic responses. We considered a poststimulus response significant when its CUSUM deflection exceeded the error box limits, an approach previously validated using in vitro brain slice recordings (*58*). For such events, we defined response latency as the earliest inflection point marking the onset of the CUSUM deflection. For inhibitory responses derived from PSF analyses, we determined latency from the last sustained discharge preceding firing cessation (*58*). This conservative criterion ensured that we only included robust inhibitory responses in subsequent analyses (*60*). We excluded motor units exhibiting weak or non-significant inhibition where CUSUM amplitudes remained below the error box threshold.

#### Statistical analyses

Statistical analyses were performed in R (version 4.3.2). Computational simulations and post-processing were conducted in Python (version 3.11) within Jupyter notebooks. Graphs were generated using Python and GraphPad Prism (version 10.0; GraphPad Software, USA), and schematic illustrations were created with BioRender. Statistical significance was set at P < 0.05 for all analyses.

#### Experimental Analysis

First, we performed separate linear regression analyses for each side (ipsilateral and contralateral) to model inhibition duration as a function of motor unit firing rate at the group level. We quantified the strength of the association between these variables using the coefficient of determination (*61*, *62*).

To examine the relationship between inhibition duration and motor unit discharge rate at the experimental level while accounting for circuit laterality, we fitted a linear mixed effects model with inhibition duration as the dependent variable. We included motor unit firing rate and side (ipsilateral vs contralateral) as fixed effects, together with their interaction, to test whether the association between firing rate and inhibition duration differed between circuits.

We included random intercepts for motor unit and participant to account for repeated measurements within motor units and the hierarchical structure of the data across participants. We mean-centred all continuous covariates prior to model fitting. We fitted the model assuming normally distributed residuals, using restricted maximum likelihood estimation with the bobyqa optimiser. We estimated degrees of freedom using the Satterthwaite approximation and used Wald confidence intervals for fixed effect estimates. We applied the same statistical approach to inhibition latency, using latency as the dependent variable in place of inhibition duration.

We compared mean motor unit firing rate between sides using a paired t-test, with side treated as a within-subject factor. In addition, we compared vastus lateralis (VL) M-wave amplitude, expressed as a percentage of M_max_, between sides using a paired t-test to ensure that firing rates and stimulation conditions were comparable across sides.

To determine whether inhibition duration was modulated by contraction intensity independently of motor unit discharge rate, we fitted a multiple linear regression model with inhibition duration as the dependent variable. Force output was included as the primary predictor, while firing rate was decomposed into between– and within-motor unit components and included as covariates. The between-motor unit component reflected differences in average discharge rate across motor units, whereas the within-motor unit component represented deviations from each motor unit’s mean discharge rate across force levels.

To further investigate whether motor unit recruitment threshold influenced inhibition duration, we fitted a linear mixed-effects model with inhibition duration as the dependent variable. Motor unit recruitment threshold and stimulation side (ipsilateral vs. contralateral), along with their interaction, were included as fixed effects to determine whether the relationship between recruitment threshold and inhibition duration differed between circuits. Random intercepts for both motor unit and participant were included to account for repeated measurements within motor units and for the hierarchical structure of the dataset across participants.

#### In silico experiment analysis

For the in-silico simulations, we quantified inhibitory responses for each MN using the same PSTH and PSF procedures applied to the experimental data. To enable direct comparison between experimental and simulated data, we used experimentally measured motor unit firing rates as inputs to all simulated linear models and evaluated predicted inhibition durations at these experimental firing rates. For each participant, we evaluated all candidate simulated linear models derived from the simulation library, which spanned different combinations of inhibitory amplitude and time constant parameters.

We based model selection on the minimization of the mean squared error (MSE) between experimentally observed inhibition durations and those predicted by each simulated model. Because the mean squared error corresponds to the residual term of the coefficient of determination, this criterion ensured that the selected simulated linear model minimized overall deviation from experimental data while preserving the dependence of inhibition duration on firing rate.

In addition, we computed a gain metric (pseudo r²) relative to a null model, where we defined the null model as the mean of the experimental inhibition durations. Positive values for this gain indicate improved predictive performance relative to the null model. To provide a comparative metric when illustrating the optimized simulated and experimental linear models for ipsilateral and contralateral recurrent inhibitions, we quantified effect size using Hedges’ g. Specifically, we used Hedges’ g to compare the distributions of squared residuals obtained from simulated model predictions and experimental linear fits.

To quantify the independent contributions of MN size, firing rate, and circuit laterality to inhibition duration, we fitted a linear mixed effects model with inhibition duration as the dependent variable. We included soma capacitance as a fixed effect indexing MN size. We decomposed motor unit firing rate into two components: the MN specific mean firing rate across all observations, capturing between MN differences at the baseline discharge level, and within MN deviations from this mean, capturing condition dependent fluctuations in firing rate within individual MNs.

We included these two components as separate fixed effects. We also included circuit laterality (side) as a fixed factor, together with its interaction with soma capacitance, to test whether the relationship between MN size and inhibition duration differed across circuits. We included MN identity as a random intercept to account for repeated measurements within MNs. We mean-centred all continuous covariates prior to model fitting. We fitted the model assuming normally distributed residuals using restricted maximum likelihood estimation. We estimated degrees of freedom using the Satterthwaite approximation and used Wald confidence intervals for fixed effect estimates.

## Author contributions

Writing—original draft: J.C, D.G and S.B. Conceptualization: J.C., D.G., and S.B. Investigation: J.C. and D.G. Methodology: J.C., D.G., and S.B. Funding acquisition: S.B. Data curation: J.C. and D.G. Validation: J.C., D.G., and S.B. Supervision: S.B. Formal analysis: J.C. and D.G. Software and modeling: J.C. and D.G. Project administration: S.B.

## Data and materials availability

All data needed to evaluate the conclusions of this paper are present in the paper and/or the Supplementary Materials. No new materials were created for this work. The datasets generated and analysed during the current study will be deposited in an institutional repository and will be available from the corresponding author upon reasonable request. All custom code used for data processing, analysis, and visualization will be made publicly available through an open-source repository upon publication.

